# Whole-genome sequencing of Bantu-speakers from Angola and Mozambique reveals complex dispersal patterns and interactions throughout sub-Saharan Africa

**DOI:** 10.1101/2022.02.07.478793

**Authors:** Sam Tallman, Maria das Dores Sungo, Sílvio Saranga, Sandra Beleza

## Abstract

As the continent of origin for our species, Africa harbours the highest levels of diversity anywhere on Earth. This trove of diversity provides rich opportunities to discover unknown genomic variation and explore how human populations have moved and interacted on the continent over thousands of years. However, many regions of Africa remain under-sampled. Here we present the first collection of whole-genomes from Angola and Mozambique, enabling the construction of a high-quality reference variation catalogue including three million novel SNPs. Leveraging the power and flexibility of whole-genome sequencing data, we model the development and continuity of Bantu-population structure through time, widespread admixture involving source populations from these regions across sub-Saharan Africa, and the heterogeneous population histories of Western and Eastern Bantu-speakers. In contrast to depictions of the Bantu expansion as a single, continuous macro-event, we recover evidence of admixture among distinct Bantu-speaking groups in South-Eastern Africa and bring together concordant patterns from linguistics and archaeology to paint a more complex picture of Iron Age migrations into the region. Moreover, we generate reference panels that better represents the complete diversity of African populations involved in the Atlantic slave trade, improving imputation accuracy in African Americans and Brazilians over the 1000 Genomes Project. This study fills important gaps in the current record of global genetic diversity and informs on the most significant demographic events in the recent history of Africa. We anticipate that our collection of genomes will form the foundation for future genomic healthcare initiatives involving under-represented communities in Angola and Mozambique.

## Main

With over 300 million speakers (5% of the global population) spanning a region of sub-Saharan Africa 10 million km^2^, Bantu represents one of the world’s largest language groups. This vast distribution has been largely attributed to the Bantu expansion, a succession of dispersals originating in the inland Savannahs of Central-West Africa some 6,000–5,000 years before the present-day (BP)^[1][2][3]^ and spanning the Iron Age (2,200–1,000 BP) likely mediated by the development of agriculture^[4][5]^ and periods of habitat change^[6][7]^. Historical records (www.slavevoyages.org) show that Bantu-speaking communities were also heavily affected by the forced export of peoples to the Americas during the Atlantic slave trade, contributing over half of all slaves to have disembarked across the New World. Despite this central role in the histories of both Africa and the Americas, Bantu-speaking communities remain underrepresented in human genomics research.

Today, advances in next-generation sequencing technologies have begun to facilitate the curation of whole-genome sequencing (WGS) data representing the full spectrum of variation across diverse human populations^[8][9]^. Such endeavours are critical next steps towards understanding how genetic diversity is structured globally and providing reference variation catalogues for a broad range of medical genetics initiatives^[10][11]^. Although Bantu-speaking communities have been involved in several WGS projects to date, such as The 1000 Genomes Project (1000G)^[12]^, African Genome Variation Project (AGVP)^[13]^, Ugandan Genome Resource (UG2G)^[14]^ and The H3Africa Initiative^[15]^, gaps remain including a scarcity of data from populations on the edge of the Bantu expansion such as those from Angola and Mozambique.

Prior analyses of autosomal SNP array^[16][17]^ and linguistic data^[18]^ from Angola and Mozambique have proved crucial in forming our understanding of major dispersal routes undertaken during the Bantu expansion. This includes favouring the so-called ‘late-split’ model concerning the diversification of Western and Eastern Bantu languages. However, archaeological^[19][20]^ and linguistic data^[21]^ have hinted at additional, more complex patterns — including successive dispersals and subsequent interactions among distinct Bantu communities in South-East Africa, suggesting our understanding of migrations into and out of Angola and Mozambique remains incomplete.

Furthermore, as former Portuguese colonies, Angola and Mozambique are recorded as being the origin of over 5 million and 500,000 slaves to have crossed the Atlantic respectively from 1526–1875 (www.slavevoyages.org). With limited genomic data available from both regions, current reference variation panels lack the complete diversity of parental African populations that have contributed significant ancestry to populations throughout the Americas^[17][22]^ potentially leading to asymmetries in our ability to describe and impute variation in these populations for analysis in Genome-Wide Association Studies (GWAS)^[23]^.

To support in the continued discovery and cataloguing of genomic variation in human populations and to further our understanding of the Bantu expansion, we sequenced the genomes of 300 individuals from Cabinda, a northern exclave of Angola, and 50 individuals from Maputo, Mozambique to a mean coverage of 12*X*. Utilizing the power and flexibility of whole-genome sequencing data, we discover rare variation, fine-scale population structure, and heterogeneous population histories of Angolans and Mozambicans. Moreover, in agreement with archaeological^[19][20]^ and linguistic data^[21]^, we perform haplotype-and model-based analyses that uncover evidence that dispersal patterns involving Bantu-speaking populations in sub-Saharan Africa were likely more complex than previously described. Finally, we generate a reference variation panel that better represents the diversity of African populations involved in the Atlantic slave trade. Overall, this dataset represents a timely addition to the growing number of whole-genome sequences from Africa, generates new insights into the history of Bantu-speaking migrant communities, and takes another step towards ensuring the potential benefits of genomics extends to all parts of the globe.

## Results

### 1 A novel collection of genomes from Angola and Mozambique uncovers rare variation

Genomic DNA was extracted using saliva samples collected with informed consent from 300 Angolans and 50 Mozambicans and sequenced using the Illumina HiSeq X™ platform to an average autosomal read depth of approximately 12*X* (Table 1) and labelled using self-reported parental and grand-parental languages (Supplementary Table 1). After sample processing, variant calling, and quality-control we identified 33.1 million total variants, including 29.9 million SNPs and 3.2 million short INDELs, with an average of 4.1 million autosomal SNPs per sampled genome (Table 1). Approximately three million SNPs were novel when compared to the dbSNP150 (https://ftp.ncbi.nih.gov/snp/), 95% of which were singletons. Modest differences in genomic diversity between collection sites were apparent, with individuals from Angolan populations showing an increased average heterozygosity compared to those from Mozambique (Wilcoxon p=6.26^-6^).

**Table 1.** Summary of newly sequenced individuals and autosomal genomic variation in Angola and Mozambique Other includes Baluba (n=1), Lingala (n=1), Lunda (n=1) and individuals with mixed grand parental languages. ± shows one standard deviation.

| Population | Angola | Mozambique |
| --- | --- | --- |
| Language | Kikongo<br>(n=238),<br>Kimbundu<br>(n=14),<br>Ovimbundu<br>(n=15),<br>Ovambo<br>(n=2),<br>Chokwe<br>(n=2), Other<br>(n=26) | Twa-Ronga<br>(n=18), Chopi<br>(n=13),<br>Makua-Lomwe<br>(n=14), Sena<br>(n=3), Shona<br>(n=1), Other<br>(n=1) |
| Total samples (post QC) | 298 | 50 |
| Mean coverage (X) | 11.56 ± 1.51 | 12.07 ± 1.40 |
| Mean reads mapped hg19/GRCh37 (%) | 90.19 ± 6.83 | 93.94 ± 2.86 |
| Total SNPs | 27,116,464 | 17,064,063 |
| Total INDELs ;50bp | 2,964,806 | 1,840,852 |
| Nonsynonymous | 134,936 | 67,349 |
| Synonymous | 112,122 | 63,876 |
| Downstream | 273,729 | 163,500 |
| Upstream | 257,040 | 163,500 |
| 5'UTR | 80,086 | 448,98 |
| 3'UTR | 327,251 | 184,218 |
| Intronic | 11,548,432 | 6,774,720 |
| Intergenic | 14,662,426 | 8,828,426 |
| Splicing | 2,509 | 1,066 |
| ncRNA | 3,950,561 | 2,347,461 |

When compared with variation generated as part of the 1000G^[13]^ and the AGVP^[14]^, 24% and 7% of autosomal SNPs segregating within Angola and Mozambique respectively were population specific. Of the 6.8 million total dataset-specific SNPs, 64% were singletons and 85% were rare (minor allele frequency (MAF) < 0.05). Upon examining shared *f2* alleles^[24]^ which appear exactly twice between individuals from Angola or Mozambique and populations from the 1000G and the AGVP, we find the largest proportion of rare variants in these newly sequenced populations are shared with other Bantu-speaking groups, with notable yet reduced sharing with other non-Bantu speaking West African and African-derived populations, especially those from the Americas (Supplementary Table 3).

### 2 The emergence and continuity of Bantu population structure in Angola and Mozambique in a pan-African context

To investigate population structure and diversity across Angola and Mozambique in a global context, we merge unrelated individuals from our newly sequenced collection with sequenced individuals from the 1000G and the AGVP (Supplementary Table 2). After removing individuals from Angola and Mozambique with evidence of recent European ancestors (as estimated by ADMIXTURE^[25]^) (Supplementary Table 1), we calculate F_ST_^[26]^ to estimate genetic differentiation between each population pair. Consistent with a demic diffusion of Bantu languages across Africa, we find low genetic differentiation (F_ST_ ≤ 0.015) amongst all Niger-Congo populations (including Bantu speakers and non-Bantu speaking West Africans) (Supplementary Table 4), with pair-wise F_ST_ decaying with geographic proximity (Spearman’s rho = 0.83, p = 2.69^-12^).

PCA^[27]^ performed on Niger-Congo speakers in this WGS dataset captures population structure following an isolation-by-distance pattern across the continent, with PC1 separating East from West (Figure 1b). Procrustes analysis supports these qualitative observations, showing a strong correlation between genetics and geography (Procrustes similarity t0 = 0.84, p < 10^-5^). Even within Mozambique, individuals form a cline that separates ethnolinguistic groups commonly found in the North (Makua-Lomwe) and South (Tswa-Ronga, Chopi) of the country. To further examine this apparent structure, we calculate *f4* statistics^[28]^ in the form *f4(Mozambique (South), Mozambique (North); X, Chimp)* where *X* is any other population in our WGS dataset. Here, the only significant (Z = 9.3) statistic tests clade structure compared with the AGVP Zulu, with a positive value of *f4* indicating increased allele sharing between the Zulu and speakers of languages commonly found in the South of Mozambique (Supplementary Table 5).

**Figure 1.**
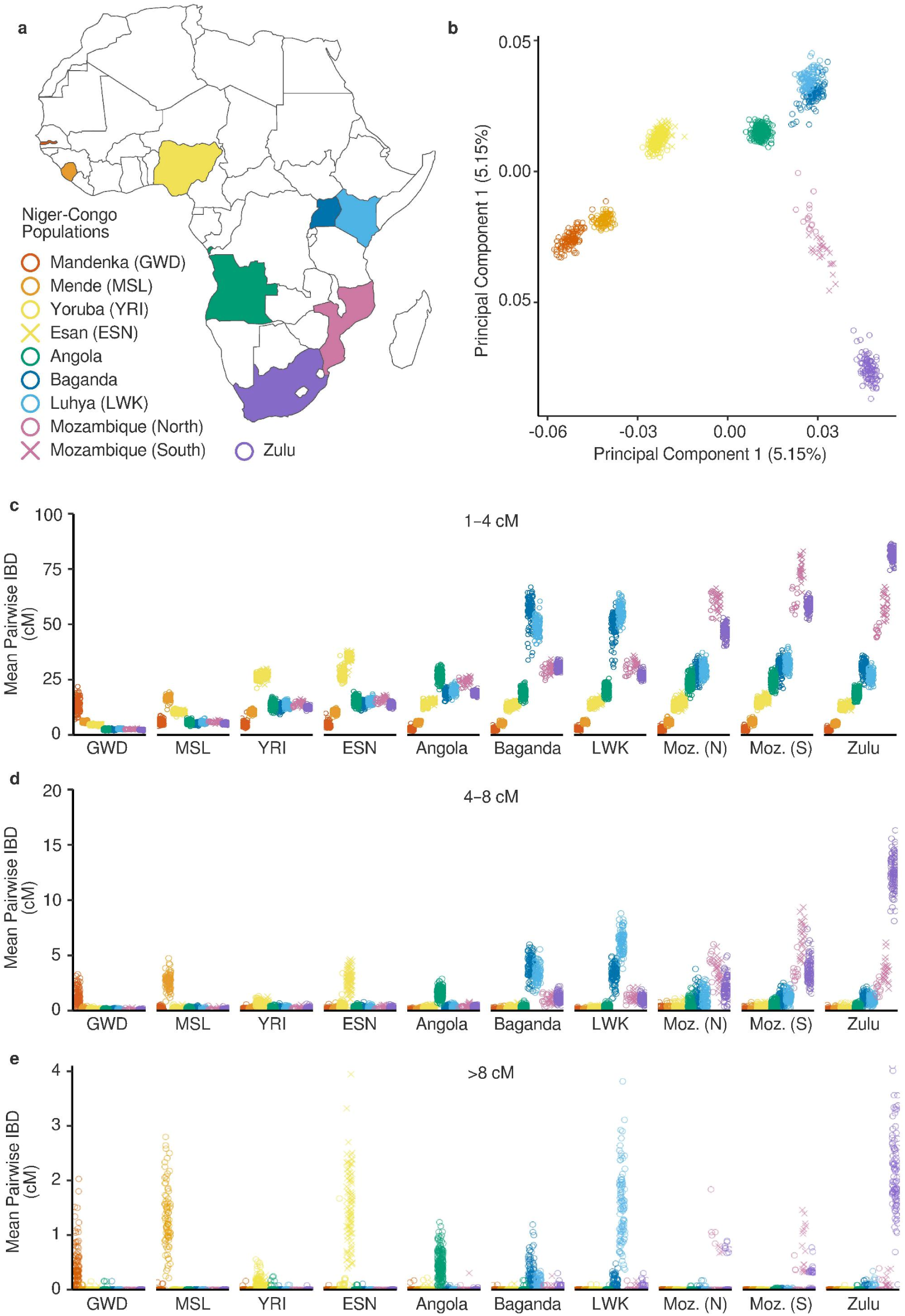
Population structure of sequenced Niger-Congo populations. (a) Map denoting colour and shape corresponding to Niger-Congo populations present in a merged dataset consisting of newly sequenced Angolans and Mozambicans, the 1000G and the AGVP (Supplementary Table 2). (b) Top two principal components of PCA of the merged dataset. (c) Average cumulative length of IBD haplotypes between 1 and 4cM that individuals share with another individual from each population in the merged dataset. (d) As in c, but for IBD haplotypes between 4 and 8 cM. (e) As in c/d but for IBD haplotypes greater than 8cM. Populations are ordered by geographic distance from The Gambia. Angola includes all individuals with a single familial language sampled from Angola. Mozambique (North) and Mozambique (South) labels were determined using genetic variation patterns and include Tswa-Ronga and Chopi or Makua-Lomwe speakers respectively. GWD, Mandenka from The Gambia; MSL, Mende from Sierra Leone, YRI; Yoruba from Nigeria; ESN, Esan from Nigeria; Angolans; LWK, Luhya from Kenya; Baganda from Uganda, Moz. (N), Northern language speakers from Mozambique; Moz. (S), Southern language speakers from Mozambique; Zulu from South Africa.

Examining Identical-By-Descent (IBD) haplotypes^[29]^ shared across Niger-Congo populations within the dataset, we observe geographic stratification of recent shared ancestry (Figure 1c/d/e). Longer, more recent haplotypes (>8cM) (Figure 1e) are shared almost exclusively within populations, consistent with more restricted population movements in recent centuries. For Angolans and non-Bantu speakers this is also true of intermediate haplotypes (4–8cM) (Figure 1d). However, Eastern (AGVP Baganda, 1000G LWK) and South-Eastern Bantu-speakers (Mozambicans, AGVP Zulu) still share ancestors in this period, illustrating their recent common histories.When focusing on shorter, more ancient haplotypes estimated across populations (1–4cM) (Figure 1c), considerable shared ancestry across all Bantu-speaking populations is observed.

Examining more ancient IBD (1–4cM) sharing within populations (Figure 1c), clear differences between regions become apparent. Bantu-speaking groups from East Africa share higher mean pairwise IBD than populations from further West (East = 64cM, West = 24cM, p < 1x10^-5^). This pattern is similarly observed when analysing short Runs-Of-Homozygosity (ROH)^[30]^ (Supplementary Figure 1). Among Mozambicans, Southern language speakers share higher mean pairwise IBD than Northern language speakers (Mozambique (South) = 75cM, Mozambique (North) = 60cM, p < 1x10^-5^), corroborating the recent discovery of North-to-South serial founder events in the genetic history of the region^[17]^. Overall, we infer a strong correlation between within-population IBD sharing and geographic distance from The Gambia (Spearman’s rho = 0.9, p = 8.8x10^-4^), indicative of a progressive reduction in genetic diversity associated with the macro-scale direction of Niger-Congo migration out of West Africa and subsequently into East and South-East Africa^[2]^.

To further explore the fine-scale population structure of Angola and Mozambique alongside neighbouring groups, we merged our extended WGS dataset with a curated selection of modern and ancient individuals genotyped at SNP sites present on the Human Origins Array (HOA)^[31][32][33][34][35][36][37][38][39][40][41]^ (Supplementary Table 6).

Unsupervised, haplotype-based clustering of Angolans and Mozambicans alongside 420 individuals present in the wider dataset using fineSTRUCTURE^[42]^ recovers fine-scale regional genetic relationships between Bantu-speaking groups (Supplementary Figure 2b). Supporting our inference above (Figure 1b), we infer genetic sub-structure within Mozambique separating Southern (Tswa-Ronga, Chopi), Northern (Makua-Lomwe) and Western (Sena, Shona) language speakers, with the latter two clustering together with neighbouring Malawians^[34]^. Angolans within our dataset could be separated into clusters separating Northern (largely Kikongo), Central (Kimbundu, Ovimbundu) and Southern (Ovambo) language speakers. Together, these results show that population structure reflecting self-reported parental and grand-parental language exists among individuals co-inhabiting urban population centres in present-day Angola and Mozambique.

Analyses of contemporary Bantu-speakers in our HOA dataset in the context of aDNA collected from Iron Age farmer sites^[31][34][35][36]^ using PCA (Supplementary Figure 2a) and ADMIXTURE (Supplementary Figure 3) shows ancient individuals appear most closely related to modern populations from neighbouring regions (1,400 year old individuals from close to the Okavango delta in Botswana are a notable exception, as described in REF^[31]^). Angolans appear closest to two 250 year old individuals from Kindoki hill in neighbouring Congo^[31]^ and Mozambicans appear closest to a 700 year old individual from Pemba Island in neighbouring Tanzania^[34]^. This suggests that, despite the rapid and widespread movement of people throughout sub-Saharan Africa over the past 6,000 years^[1][2][3][13][14][16]^ (Figure 2), there has been a relative continuity of regional Bantu population structure over recent centuries.

**Figure 2.**
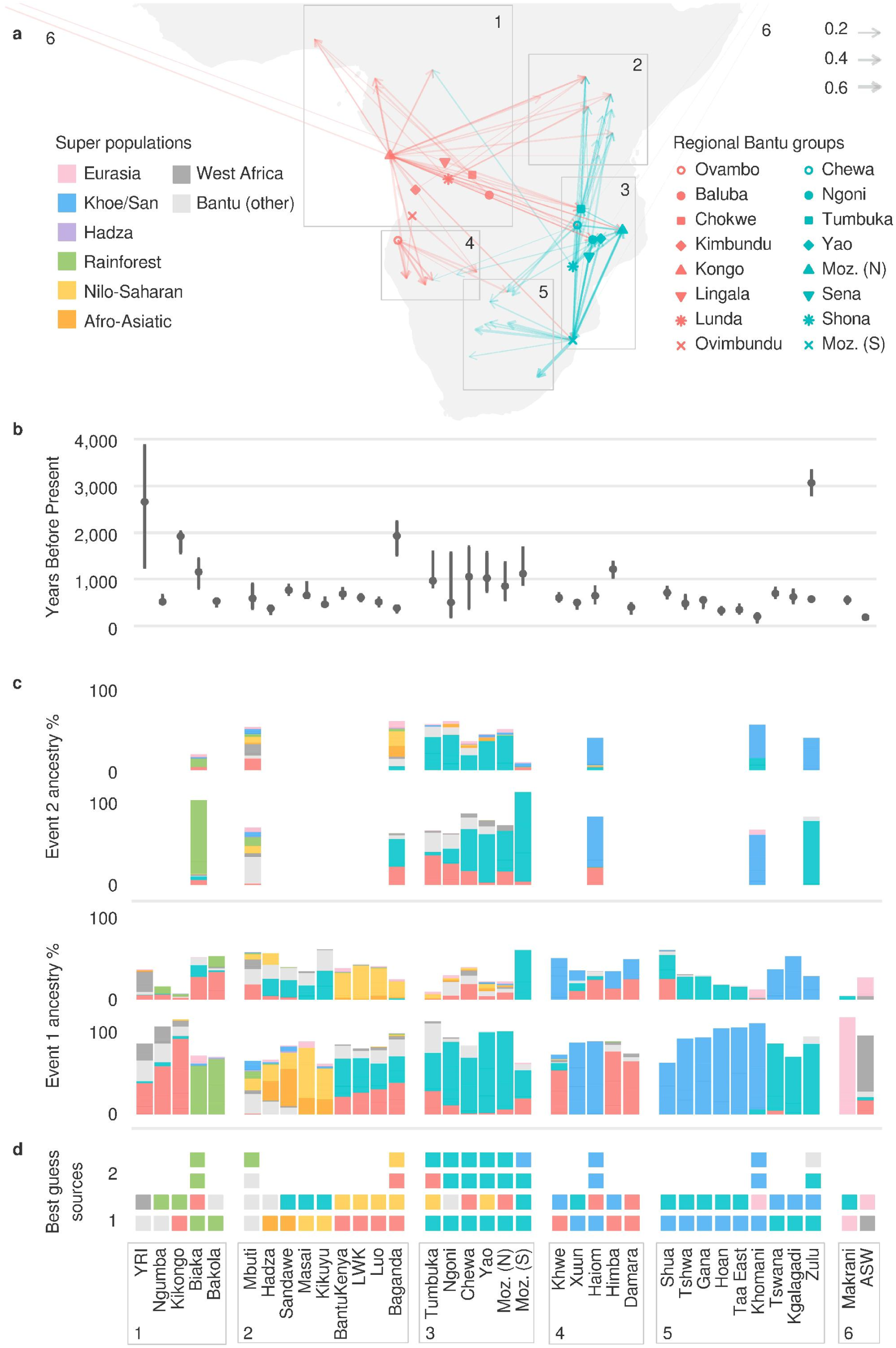
Complex admixture involving Central-West and South-Eastern Bantu-speakers over the past 4,000 years as inferred by fastGLOBETROTTER. (a) Map arrows connecting 36 African and non-African groups with evidence of admixture involving source populations at least partially (>5%) matched to Angolan (red), Mozambican, or Malawian (light blue) Bantu-speaking donor populations. The size of the arrow is proportional to the fraction of ancestry from the donor population. Coordinates for each source population were selected based on the approximate region of Africa where that language is primarily spoken. (b) Date(s) of admixture estimated using fastGLOBETROTTER. Generation time was transformed into years using 26 years per generation. (c) Modelled ancestry composition of the admixing source populations inferred during either events 1 (latest) or 2 (earliest). In instances where fastGLOBETROTTER infers a multi-way event involving more than two source populations mixing at a single date (e.g. in the case of Mozambicans and Malawians), we describe the estimated ancestry composition of the less strongly-signalled source populations under event 2. However, we note that determining the precise ancestry composition of source populations involved in these multi-way events can be difficult (see REF^[45]^). Colours represent groups merged into super populations for ease of visualisation (see Supplementary Table 7 for full results). (d) Ancestry of the best guess major (M) and minor (m) source populations involved in events 1 and 2 inferred by fastGLOBETROTTER. Mozambique (North) and Mozambique (South) labels were determined using genetic variation patterns and include Tswa-Ronga and Chopi or Makua-Lomwe speakers respectively. YRI; Yoruba from Nigeria; LWK, Luhya from Kenya; Moz. (N), Northern language speakers from Mozambique; Moz. (S), Southern language speakers from Mozambique.

### 3 Complex admixture and widespread migration into and out of Angola and Mozambique over the past 3,500 years

The genetic architecture of sub-Saharan Africa has been shaped by admixture involving Bantu migrants and autochthonous populations^[18][35][40]^. Consistent with a history involving almost complete replacement of local genetic diversity across large parts of Central Africa^[15][16][17][34}^, ADMIXTURE^[25]^ (Supplementary Figure 3) and SOURCEFIND^[43]^ (Supplementary Figure 4) analyses performed using our extended HOA dataset (Supplementary Table 6) suggests that Angolans and Mozambicans are best represented as having 98% (minimum (min) = 89%, maximum (max) = 100%, standard deviation (sd) ± 2%) and 97% (min = 92%, max = 99%, sd ± 2%) West African (Bantu-related) ancestry respectively. This is also true of neighbouring Malawian and Namibian Bantu-speakers^[34][41]^. Using fastGLOBETROTTER^[44][45]^ (Figure 2, Supplementary Table 7), we find a small 2–3% contribution from a Khoe/San source group (best represented by a 2,000 year old South African from Ballito Bay^[35]^) in Mozambique (South) derived from a single admixture event approximately 1,100 BP (95% confidence interval (CI) 900–1,650 BP), pre-dating the most recent Khoe/San admixture event observed in neighbouring South Africans (AGVP Zulu) by approximately 500 years (Figure 2b). However, we do find evidence for an earlier admixture event involving Mozambique (South) and Khoe/San groups in the Zulu at approximately 3,000 BP (CI 2,800–3,300 BP), suggesting interactions with autochthonous populations may have occurred during an earlier phase of Bantu migration into the region. We also infer admixture involving a small 3–4% Western rainforest hunter-gatherer component (best represented by the Cameroonian Bakola^[38]^) occurring at approximately 1,900 BP (CI 1,600–2,000 BP) in Angolan Kikongo speakers. Individuals who speak Central Angolan or North Mozambican languages are modelled as having as little as <1% hunter-gatherer related ancestry.

However, of the 39 sub-Saharan African populations in our HOA dataset whose range overlaps with Bantu languages, 27 (70%) had evidence introgression from a source population at least partially (>5%) matched to Angolans or Mozambicans (or closely related Malawians) (Figure 2, Supplementary Table 7). Thus, whilst admixture with autochthonous populations have had a modest impact on the gene-pool of the present-day Bantu-speaking inhabitants of Angola and Mozambique (Supplementary Figure 3/4), it is likely that the ancestors of these groups mixed extensively with populations from across sub-Saharan Africa, likely reflecting biased mating practices of Bantu-speaking agriculturalists and local hunter-gatherers (see REF^[34]^). Furthr examining the geographic distribution of these inferred admixture events recaptures an extensive history of Bantu migration into and out of Central-West and South-East Africa before, during, and after the Iron Age (Figure 2a). Admixture events involving populations with Angolan-like ancestry appear over a time transect spanning approximately 3,500 BP and an area of over 10,000,000 km^2^, consistent with massive migration across West Africa and into East Africa after the development of agriculture and the fragmentation of the Central African rainforests^[5][6][7][16][18]^. On the other hand, ancestry matched to Mozambicans and Malawians highlights the impact of later Eastern Bantu dispersals, uncovering evidence of extensive migration between this region and both South Africa/Botswana as well as the Great Lakes region to the North (Figure 2a)^[17][19][20][46]^.

Notably, in this same analysis, we also infer complex, multi-way admixture in Mozambican and Malawian populations occurring on average approximately 1,000 BP (CI 500–1,700 BP) (Figure 2, Supplementary Table 7). In accordance with archaeological^[19][20]^ and linguistic research^[21]^ suggesting that Bantu-speaking communities arrived in South-East Africa via multiple waves of migration, these admixture events likely involved multiple sources variably related to both Eastern and Western Bantu-speaking groups, in addition to the small (1-6%) contributions from autochthonous populations (Supplementary Figure 5, Supplementary Table 7). Importantly, detection of admixture events among distinct Bantu-speaking groups in South-East Africa provide further evidence against simple, continuous migration models of the Bantu expansion^[3]^.

### 4 Distinct periods of population size change, diversification, and gene flow in the population histories of Angolans and Mozambicans

Genomic data can reveal insights into ancestral demography over time and the formation of present-day populations. Free from the confounding effects of substantial recent autochthonous admixture (Figure 2), Angolans and Mozambicans are well-placed to investigate demography specific to their respective Western and Eastern Bantu ancestors. To further investigate the population size and separation histories of Angolans and Mozambicans, we use the non-parametric Multiple Sequentially Markovian Coalescent method (MSMC2)^[47]^ and genome-wide genealogies estimated using Relate^[48]^. We also investigate the demographic histories of Angolans and Mozambicans under an Approximate Bayesian Computation (ABC) framework^[49][50]^, simulating 200,000 whole-chromosomes^[51]^ (chromosome 1) with realistic, locus-specific recombination rates and error rates typical of WGS data^[52]^. This simulated data was then compared to real data across a set of summary statistics (Supplementary Table 8), selected based on their ability to inform on various demographic parameters. This includes IBD and ROH haplotype-based statistics^[31][32]^ that require whole-chromosome-scale recombination maps to accurately calculate, making them inaccessible to ABC methods until recently^[53]^.

Concerning population size histories, concordant patterns from all three methodologies are observed. Here, Mozambicans are estimated as having a lower ancestral *Ne* than Angolans (Figure 3a/b/c/e), perhaps due to a bottleneck associated with the later dispersal of Bantu-speaking communities into or within South-East Africa. This was followed by rapid growth in both populations (Figure 3a/b/d/f), likely driven by a transition to more sedentary lifestyles^[54]^.

**Figure 3.**
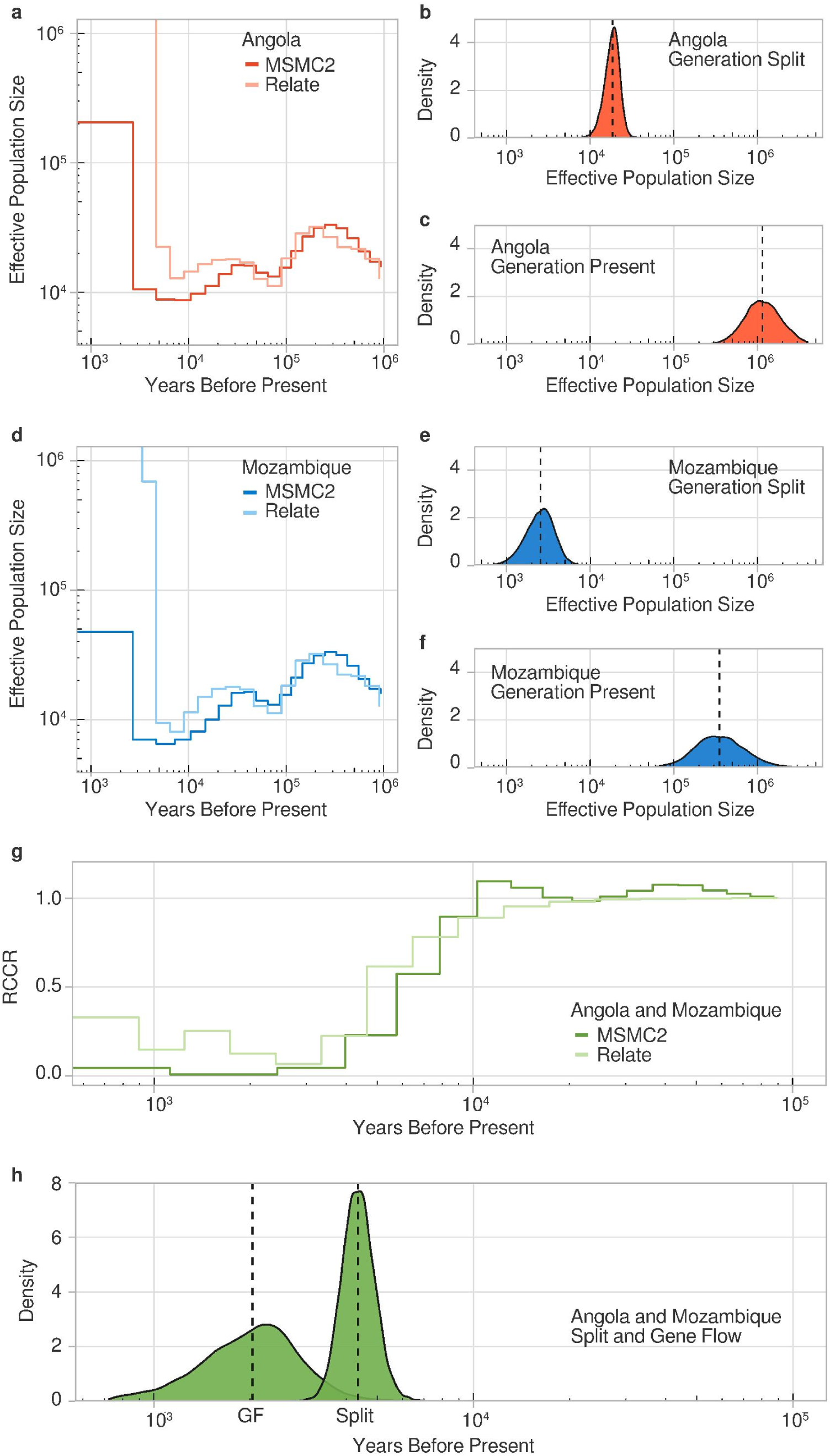
Demographic models of Angolan and Mozambican population size and separation histories. (a) Effective population size history of Angolans using within-population coalescence rates estimated using Relate with 40 genomes and MSMC2 with 4 high-coverage genomes (b) ABC posterior distribution for parameter denoting Angolan effective population size at the generation in which Angolans and Mozambicans separated (Generation Split) (c) ABC posterior distribution for parameter denoting Angolan effective population size in most recent generation (Generation Present). (d/e/f) As in a/b/c but for Mozambicans). (g) Separation history of Angolan and Mozambicans estimated using Relative Cross-Coalescence Rates (RCCR) estimated using Relate with 40 genomes from each population and MSMC2 with two high-coverage genomes from each population. Separation times were taken as the first generation going backwards-in-time in which RCCR is greater than or equal to 0.5 (h) ABC posterior distributions for parameters denoting the generation during which Angolans and Mozambicans first split (Generation Split) and the generation during which Angolans and Mozambicans subsequently exchanged migrants (Generation Gene Flow (GF)). Dashed lines represent the medians of the posterior distributions. Generation time was transformed into years using 26 years per generation. Angola includes a subset of individuals with a single familial language sampled from Angola. Mozambique includes all individuals with a single familial language sampled from Mozambique.

Using MSMC2 or Relate to determine the Relative-cross coalescence rates (RCCR), place the split between Angolans and Mozambicans at 5,500 BP and 4,300 BP respectively (generation where RCCR > 0.5, Figure 3g). As expected from the relative homogeneity of Bantu language speakers, these split time estimates are more recent than those estimated between Angolans or Mozambicans and non-Bantu speaking groups such as the Yoruba (1000G YRI), Sierra Leonian Mende (1000G MSL) and the Gambian Mandenka (1000G GWD) (Supplementary Figure 6). However, both MSMC2 and Relate assume these splits occurred without any subsequent admixture. Conversely, our ABC framework enables demography to be investigated under more complex scenarios, such as admixture involving sources with Angolan-like ancestry in the history of Mozambicans predicted using fastGLOBETROTTER (Figure 2). To select between two models of Angolan and Mozambican population splits, namely a *clean split* or a *split with gene flow* (Supplementary Figure 7) we trained an ABC random-forest classifier^[49]^ using 100,000 simulations of each model, obtaining a posterior predictive accuracy of 80%. Applying this classifier to data from our newly sequenced Angolans and Mozambicans, we infer a 95% posterior probability of the data fitting the *split with gene flow* model, further evidencing a complex separation history. Calculating posterior parameter estimates^[50]^ under the accepted model (Supplementary Table 9/10) places the initial Angolan and Mozambican split at approximately 4,250 BP (95% CI 3,500–5,400 BP), whilst estimating additional migration between the two populations at approximately 1,700 BP (95% CI 800–3,000 BP) (Figure 3h).

Although estimates of admixture proportions are not well constrained using our ABC approach (Supplementary Figure 8), posterior distributions suggest asymmetrical gene flow from Angolan-like groups into Mozambicans (Supplementary Table 10). These results support a model involving multiple waves of dispersal into South-East Africa resulting in admixture between distinct Bantu-speaking populations.

### 5 Increasing West African reference panel diversity using Angolan and Mozambican genomes improves imputation accuracy in African Americans and Brazilians

Forced migrations from Angola and Mozambique during the Atlantic slave trade had a widespread impact on the ancestry many of present-day inhabitants of the Americas^[17][23]^. However, due to the limited availability of sequencing data, populations from these important embarkation regions are not represented in current WGS reference panels. Historical records (www.slavevoyages.org) suggest that 96% of all slaves to arrive in South-East Brazil left from ports located in Angola and Mozambique. Moreover, 25% of all slaves to arrive in the USA originated from these same regions in addition to those that disembarked from ports across much of coastal West Africa.

Studies have consistently demonstrated the value of including population-specific reference genomes alongside a more cosmopolitan collection of samples when imputing unobserved genotypes in target datasets^[13][14]^. Unsurprisingly, we observe a substantial improvement in imputation accuracy among Angolans and Mozambicans when utilizing our novel collection of whole-genome sequences in reference panels (Supplementary Figure 9). However, it is less clear whether increasing the diversity of parental African populations in reference panels would result in improvements in imputation accuracy among admixed populations from the Americas. To test this, we used two target datasets: The 23&Me African American Sequencing Project (AASP)^[55]^, including 2,303 individuals from the USA and the *Saúde Bem Estar e Envelhecimento* project (SABE)^[56]^, including 1,171 individuals from São Paulo, Brazil. ADMIXTURE clustering shows an average of 72% West African ancestry in individuals from the AASP (min = 30%, max = 100%, sd ± 8%) and 11% in individuals from SABE (min = 0%, max = 98%, sd ± 21%) (Supplementary Figure 10). After masking genotypes in each dataset other than at reference coordinates present in the Illumina Omni 2.5 Array, we imputed genotypes using either the 1000G reference panel or a combined reference panel including the 1000G and our novel collection of genomes from Angola and Mozambique. In accordance with our understanding of slave origins, the addition of Mozambicans and Angolans to the 1000G reference panel resulted in improvement in imputation accuracy in both individuals from the USA and Brazil (Figure 4), especially for rare (0.01 < MAF ≤ 0.05: AASP 1000G R^2^ > 0.77, AASP 1000G + AM R^2^ > 0.80; SABE 1000G R^2^ > 0.72, SABE 1000G + AM R^2^ > 0.76) and very rare (MAF ≤ 0.01: AASP 1000G R^2^ > 0.22, AASP 1000G + AM R^2^ 0.26; SABE 1000G R^2^ > 0.23, SABE 1000G + AM R^2^ > 0.30) variants.

**Figure 4.**
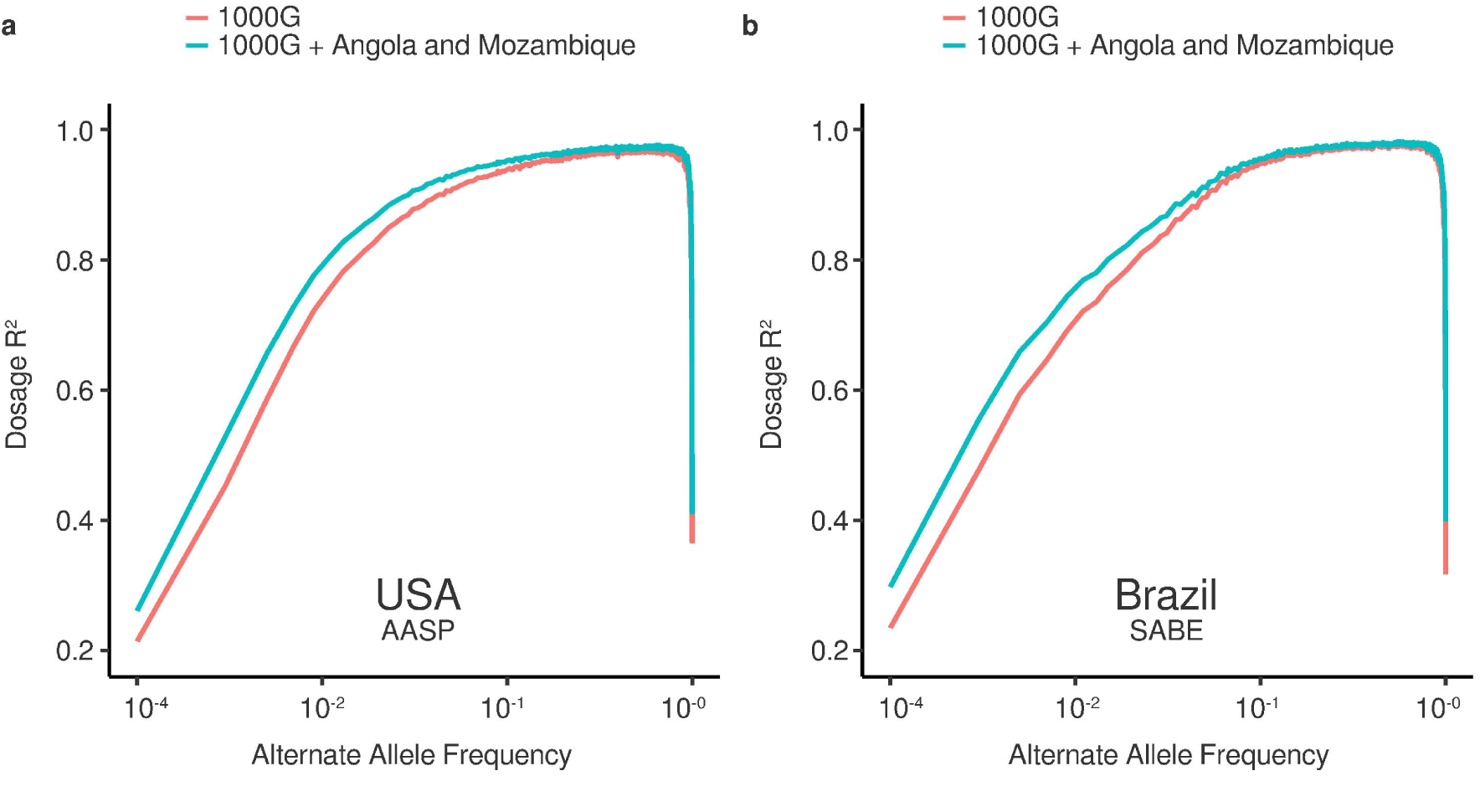
Imputation accuracy among Brazilians and African Americans from the USA after combining Angolans and Mozambicans with the 1000 Genomes Project (1000G) reference panel (a) Dosage R^2^ (Pearson’s squared correlation coefficient) of called genotype vs genotypes imputed into the USA AASP^[55]^ using either the 1000G reference panel or a merged reference panel including the 1000G and newly sequenced genomes from Angola and Mozambique as a function of alternate allele frequency. (b) As in a but for the Brazilian SABE^[56]^ dataset.

## Discussion

Analysing genomic data from under-represented human populations has the potential to shed light on major events in our species’ history and to fill important gaps in the current record of global diversity. Here, we present the first collection of whole-genomes from Angola and Mozambique, a critical step towards furthering human genomics research involving communities from both regions, expanding the coverage of catalogued genetic variation from Bantu-speakers, and potentially improving the imputation of African ancestry and African-derived ancestry in the Americas (Figure 5).

Supporting earlier discoveries^[57]^, we observe heterogeneity in the demographic histories of Angolans and Mozambicans prior to recent, independent explosions in population size (Figure 1c/d/e, Figure 3a/b/c/d/e/f). These findings provide a genetic parallel with the archaeological record, indicating a slower, more diffuse migration through the Central African rainforest into the region surrounding the Congo basin^[6][58]^ and a comparatively rapid movement of Bantu-speaking populations into and across East Africa^[5][46][59]^. Interestingly, patterns revealed by analysing ancient IBD haplotypes from sequenced West African populations (Figure 1d) suggest that serial-founder events may have accompanied the expansion of Niger-Congo languages across the continent (Figure 1c), a pattern previously unseen when analysing Y-chromosomal markers^[21]^. Analyses including a wider range of sequenced populations are warranted to test whether these differences in ancestral population sizes are broadly conserved across geographic regions.

Furthermore, we recover estimates of a genetic split between Angolans and Mozambicans beginning 3,500–5,400 BP (Figure 3e/f). This correlates with the first archaeological evidence of sedentary settlements outside of Western Cameroon ^[601^ and the emergence of familiar savannah corridors some 4,000 BP during the Mid-Holocene Rainforest Crisis, suggested to have driven early diversification of Bantu communities around the Sangha-Mbam confluence area^[7]^. Split times estimated using genomic data are less accurate in recent generations and rely on accurate mutation rate^[61]^ and generation intervals^[62]^ and thus should be treated with caution. However, these estimates do predate the earliest evidence of Bantu-associated artefacts in Eastern side of Lake Tanganyika approximately 3,000 BP^[63]^ and Mozambique approximately 2,450 BP^[64]^ as well as our earliest inferred date of admixture involving South-Eastern Bantu-speaking groups approximately 3,000 BP (Figure 2) and thus appear chronologically plausible if present-day Mozambicans trace their ancestry back to these early inhabitants.

Indeed, following on from these recent reports^[15][16][17][34}^ we find that the migrant ancestors of Angolans and Mozambicans replaced local inhabitants after their expansion across the continent, observing limited post-settlement admixture (Supplementary Figure 3/4). However, we find evidence for complex and widespread admixture involving Bantu-speaking groups from these regions (Figure 2). Perhaps most notably, we find evidence for admixture involving Western and Eastern Bantu-speaking source groups in South-East Africa approximately 800–1,700 BP (Figure 2, Figure 3h) indicative of multiple waves of dispersal into the region. Linguistic distances between Eastern and Western Bantu languages in Angola and Mozambique are smaller than those in non-adjacent regions^[21]^, suggesting that this period of admixture coincided with cultural exchange. Moreover, this multi-wave scenario resembles archaeological models that have postulated the convergence of independent Bantu dispersals in South-East Africa associated with distinct ceramic traditions, namely the Kwale and Kalundu, predicted to have connected this region to the Great Lakes to the North and Angola to the West respectively^[19]^. Interestingly, this proposed admixture also aligns well with the replacement of early farmer sites in the recorded archaeology of Mozambique by sites containing the complete Iron Age package around 1,200–1,600 BP^[20]^ and post-dates the large-scale movements of Bantu-speaking communities predicted to have been facilitated by the Late Holocene Rainforest Crisis and the opening of the Sangha River corridor after 2,500 BP^[6][7][18]^. This perhaps suggests that habitat change and parallel technological developments were drivers of later dispersals into South-East Africa. More detailed analyses of modern and ancient Bantu-speakers from sparsely sampled regions of Central Africa (see REF^[15]^) will likely refine these findings.

In conclusion, this study contributes to the ongoing effort to describe global genetic diversity and to expand our knowledge of major events in our species history. The results presented here represent another step towards a synthesis between genetic, linguistic, and archaeological research into the Bantu expansion which has at times appeared to be in conflict^[65]^ but are now beginning to paint a more complete picture of human dispersals and interactions throughout sub-Saharan Africa^[31][66]^. As with early genetic research into the Bantu expansion, which concluded that the distribution of Bantu languages was driven largely by a demic diffusion originating in West Africa^[1][67]^, it appears likely that the complex archaeological record of South-East Africa^[19]^ was also the result of human migrations and interactions.

Finally, we hope that this data will provide a reference for future research in Angola and Mozambique, including regional studies into phenotypic variation and disease susceptibility, aiding the continued emergence of new discoveries from Africa in the genomics-era.

## Methods

### 1 Sample collection and sequencing

This project was approved by the ethics committees of the University 11^th^ of November (“Universidade 11 de Novembro”), Cabinda, Angola (REf: UoN/2016), Pedagogic University (“Universidade Pedagógica”), Maputo, Mozambique (REf: UP/2017), and the University of Leicester ethics committee (REf: 11334-sdsb1-genetics). After obtaining full participant consent, saliva samples were collected from Cabinda, Angola and Maputo, Mozambique and isolated at the University of Leicester. Isolated DNA from 300 Angolan participants and 50 Mozambicans were selected and shipped for 15X target WGS. Reads of length 150 base pairs (bp) were generated by Illumina HiSeq X™. Four individuals from Angola and Four individuals from Mozambique were additionally selected for high-coverage PCR-free 50X sequencing (used in MSMC2 analyses: Methods 14).

### 2 Processing sequencing data and variant calling

minimap2 v2.11-r797^[68]^ (mode: sx) was used to map FASTQ format-ted paired-end reads generated from each newly sequenced sample against the GRCh37 reference genome (https://www.ncbi.nlm.nih.gov/assembly/GCF_000001405.13/). Resultant CRAM files were sorted according to linear reference coordinates, and duplicate reads were marked with samtools v1.9 markdup^[69]^. Base quality-scores were re-calibrated with GATK v4.0.2.1^[70]^ BaseRecalibrator and ApplyBQSR. Read depth statistics were generated from CRAM files using mosdepth v0.2.3^[71]^ with contaminated or low-quality samples removed. CRAM files were used as input for GATK v4.0.2.1 to jointly call variants across all remaining samples using the HaplotypeCaller command. This set of samples and variants were filtered and refined as described in Supplementary Note. Gene-based annotation of SNPs was performed using ANNOVAR^[72]^ utilising the GENCODE release 31 (http://ftp.ebi.ac.uk/pub/databases/gencode/Gencode_human/release_31/) and the dbSNP150 (https://ftp.ncbi.nih.gov/snp/) aligned to hg19/GRCh37.

### 3 Dataset merging and curation

SNP genotypes generated as part of our Angolan and Mozambican call-set were combined with either (a) genotype data from 2,824 individuals sequenced and genotyped as part of The 1000 Genomes Project Phase 3 (1000G)^[12]^ and the African Genome Variation Project (AGVP)^[13]^ (Supplementary Table 2) (henceforth referred to as the WGS dataset) or (b) an additional 2,394 modern and ancient individuals from 261 populations genotyped at 594,924 autosomal SNPs present in the Human Origins Array panel (HOA) described in many previous studies^[31][32][33][34][36][37][39][40][41]^ (Supplementary Table 6) (henceforth referred to as the HOA dataset). Included in the HOA dataset were three high-coverage ancient genomes: a 2,000-year-old individual from South Africa (Ballito Bay; baa001)^[35]^, a 4,500-year-old individual from Ethiopia (Mota; GB20)^[33]^ and an 8,000-year-old individual from Cameroon (Shum Laka; I10871)^[39]^. BAM files from these ancient samples were processed and subject to diploid variant calling using the procedure outlined in Schlebusch et al.^[35]^ Supplementary Materials 4.1. Both merged datasets were also combined as required with genotypes from the panTro5 Chimpanzee reference genome (https://www-ncbi-nlm-nih-gov.ezproxy3.lib.le.ac.uk/assembly/GCF_000001515.7/) aligned to hg19/GRCh37 using chain file (https://hgdownload.soe.ucsc.edu/goldenPath/panTro5/liftOver/panTro5ToHg19.over.chain.gz) provided by the the UCSC^[73]^. In each dataset, genetic distances were set using the 1000G^[12]^ genetic map (https://github.com/joepickrell/1000-genomes-genetic-maps). Extended details of our merging procedure can be found in Supplementary Note.

### 4 *f2* alleles

Shared *f2* alleles (see REF^[24]^) between populations were estimated using a custom R script (*github link placeholder*) for 40 randomly selected samples from either Angola or Mozambique all other populations within our WGS dataset. Variants within low-complexity regions (https://github.com/lh3/varcmp/raw/master/scripts/LCRhs37d5.bed.gz) and regions of known segmental duplications (https://humanparalogy.gs.washington.edu/build37/build37.htm) were ignored.

### 5 Hudson’s F_ST_

Hudson’s F_ST_ and its standard error via block-jack-knife over 1000 markers for each population-pair (where n>10) from our WGS dataset was calculated using the average hudsons_fst command within the sci-kit allel python package (https://scikit-allel.readthedocs.io/en/stable/). SNPs in relative LD were removed using PLINK v2.00a^[74]^ with parameters --indep-pairwise 50 5 0.5 as well as sites in regions of known long-range LD^[75]^ prior to calculation.

### 6 ADMIXTURE

ADMIXTURE v1.22^[25]^ was applied to two matrices of genotypes from either our WGS dataset or a subset of individuals from our HOA dataset (Supplementary Figure 4). We first pruned the data to keep sites in approximate linkage-equilibrium using PLINK v2.00a^[74]^ with parameters --indep-pairwise 50 5 0.1 whilst also excluding sites in regions of long-range linkage disequilibrium (LD)^[75]^. Ten independent, unsupervised replicates of the software were run for values K = 2,..,12. For each value K, we retain the run with the highest log-likelihood after convergence.

### 7 Principal Components Analysis (PCA)

We performed PCA on genotype matrices generated from either our WGS dataset or HOA dataset using the smartpca program (outlierremoval: NO) from the EIGENSOFT v7.2.1 tool-suite^[27]^. LD correlated SNPs were removed *a priori* using PLINK v2.00a^[74]^ with parameters --indep-pairwise 50 5 0.8. SNPs is regions of known long-range LD^[75]^ were also removed. For PCA performed on the HOA dataset, a subset of modern Bantu-speaking individuals (Supplementary Figure 2a) were used to construct eigenvectors and least-squares projection (lqproject: YES) was performed to overlay data from ancient Bantu samples, with shrinkmode: YES used to mitigate errors. For PCA performed on the WGS dataset, we restricted the data to only include Niger-Congo speakers (Figure 1b) and further performed Procrustes analysis comparing matrices of population-specific latitude and longitude coordinates and the corresponding top two principal components using the *vegan* R package^[76]^ with significance estimated via 100,000 permutations of the PROTEST command.

### 8 *f4s*tatistics

We calculate *f4* using the R package admixr^[77]^ (a port of the software ADMIXTOOLS^[28]^) across all possible four-population arrangements of Niger-Congo speakers in our WGS dataset with panTro5 set as the outgroup. Significance (Z-scores) and standard errors are estimated using a weighted block-jacknife over segments of 5-centimorgans (cM).

### 9 Identity by Descent (IBD)

IBD haplotypes were estimated across all Niger-Congo speakers in our WGS dataset using GERMLINE^[29]^ with all genotypes jointly phased *a priori* using SHAPEITv2^[78]^. We removed gaps between IBD segments that have at most one discordant homozygote and are <0.6 cM in length as well as IBD segments in regions of low SNP-density. After calculation, IBD segments <1cM were filtered out. Significant differences in the mean pairwise IBD between groups (t) was calculated by resampling groups with replacement, re-calculating the mean (t0), and treating the proportion 9,999 permutations of this procedure wherein t0 > t as our p-value.

### 10 Runs of Homozygosity (ROH)

ROH were estimated across all Niger-Congo speakers in our WGS dataset using PLINK 1.9^[74]^ with parameters --homozyg-snp 50 --homozyg-kb 300 --homozyg-density 50 --homozyg-gap 1000 --homozyg-window-snp 50 --homozyg-window-threshold 0.05. We retain only segments between 0.8 and 2.5 Mb.

### 12 CHROMOPAINTER

After phasing genotypes using SHAPEITv2^[78]^ and following the stepwise procedure to estimate global mutation/emission (-M) and switch rate (-n) parameters as outlined in previous studies^[40][79][80]^ CHROMOPAINTERv2^[42]^ was run on all diploid individuals within our HOA dataset (Supplementary Table 6) using two distinct donor-recipient population configurations: (a) all individuals and/or populations in the dataset are included as both recipients and donors of shared haplotypes (henceforth referred to as the *‘all-copying model’*) and (b) as in (a) but with all Bantu-speaking groups other than the Cameroonian Lemande (used to describe Bantu-related ancestry in previous studies^[36][39]^) excluded as donor populations (henceforth referred to as the *‘no-Bantu-copying model’*).

### 13 fineSTRUCTURE

fineSTRUCTURE v2.1.3^[42]^ was run on a subset of populations in our HOA dataset (Supplementary Figure 2b) using the chunk counts sharing matrix output from the *all-copying model,* sampling cluster assignments every 10^5^ iterations across 10^6^ total MCMC iterations after 10^6^ burn-in steps. All other individuals were fixed as super-populations (Supplementary Table 6). We next performed an additional 10^5^ hill-climbing iterations, starting from the MCMC sample with highest posterior probability. This resulted in a classification of 43 clusters that were subsequently merged into a tree using fineSTRUCTURE’s greedy algorithm.

### 14 SOURCEFIND

For a subset of individuals in our HOA dataset (Supplementary Figure 5), we used the sample-size correcting, Bayesian mixture modelling approach employed by SOURCEFINDv2^[43]^ to identify the relative proportions of ancestry that each individual shares with each given donor group using the chunk lengths sharing matrix output from the *no-Bantu-copying model*. All donor populations were provided as possible surrogate groups. The truncated Poisson prior on the number of surrogate populations that contribute ancestry to each target individual to was fixed to four, allowing eight total groups to contribute some proportion of ancestry at each MCMC iterations. We ran 200,000 total MCMC iterations and 50,000 burn-in steps, sampling mixture coefficients every 5,000 iterations. Final ancestry proportions are reported as the average of these mixture coefficients across all posterior samples.

### 15 fastGLOBETROTTER

For every population in our HOA dataset (Supplementary Table 6), we used fastGLOBETROTTER to estimate admixture^[44][45]^. Specifically, fastGLOBETROTTER requires both chunk lengths sharing matrices and individual painting sample files as inputs. Thus, to avoid self-copying between individuals within their own population, which may mask signatures of recent admixture^[80]^, we use painting sample files generated for each target population using all given donor groups except those from the target population. To avoid such occurrences in the *all-copying model*, which explicitly allows for all individuals to copy from all others, we re-ran CHROMOPAINTERv2 for each population whilst providing all other populations as donors -excluding very closely related individuals from different ethnolinguistic groups that cluster together using fineSTRUCTURE (Supplementary Figure 2b) -and using the same global mutation/emission and switch rate parameters as estimated previously. Chunk length sharing matrices were generated using the original CHROMOPAINTERv2 run from the *all-Bantu-copying model.* For each fastGLOBETROTTER run, we performed five iterations of the algorithm, generating p-values and 95% confidence intervals using bootstrap re-sampling of populations over 100 replicates. As recommended, we report results with the null.ind parameter set to 1 to avoid inference based on spurious decay signals not attributable to genuine admixture. To gain further insights into the specific donor populations being used as distinct admixing sources, we performed a visual inspection of the coancestry curves generated for each population with significant evidence of admixture and an R^2^ > 0.2.

### 16 MSMC2

MSMC2^[47]^ was used estimate within population (four individuals per population) and cross-population (two individuals per population) coalescence rates using high-coverage (50X) genomes representing individuals from both Angola and Mozambique as well as Niger-Congo speaking populations (Yoruba, Mende, Mandenka, BantuKenya, BantuTswana) sequenced as part of the Simmons Genome Diversity Project (SGDP^[37]^). SNP calls and coverage masks for each genome were generated directly from sample-specific BAM files using the *bamCaller.py* script from the MSMC GitHub repository (https://github.com/stschiff/msmc-tools) and subsequently phased using SHAPEITv2^[78]^ alongside 1000G^[12]^ reference panel (https://mathgen.stats.ox.ac.uk/impute/1000GP_Phase3.html). Following recommendations, we use sample-specific masks to exclude genotypes present regions of low coverage relative to the genome-wide average with coverage statistics generated using mosdepth^[71]^. We also exclude genotypes across all samples using Heng Li’s universal mask^[37]^. The mutation rate used to scale time was 1.25 x 10^-8^ per base-pair per-generation^[61]^.

### 17 Relate

Relate v1.1^[48]^, was used estimate genome-wide genealogies using 40 randomly sub-sampled, unrelated Angolan genomes and 40 randomly subsampled, unrelated Mozambican genomes. Genotypes were phased using SHAPEITv2^[78]^ and filtered at sites marked as “not passing” in the 1000G accessible genome pilot mask (ftp://ftp.1000genomes.ebi.ac.uk/vol1/ftp/release/20130502/supporting/accessible_genome_masks/StrictMask/). The 6-EPO multiple alignment estimation of the human ancestral genome (http://ftp.1000genomes.ebi.ac.uk/vol1/ftp/phase1/analysis_results/supporting/ancestral_alignments/) was used to identify the most likely ancestral allele for each locus. Within-population and cross-population coalescence rates were calculated using the EstimatePopulationSize.sh script from the Relate Git-Hub repository (https://myersgroup.github.io/relate/).

### 18 Approximate Bayesian Computation (ABC)

We first define two simple, two-population demographic models (Supplementary Figure 7). One describes a simple clean-split and one that additionally involves gene-flow between populations after an initial split. Here populations *P_Angola_* and *P_Mozambique_* represent populations of Bantu-speakers from present-day Angola and Mozambique respectively.

1 *Clean Split.* Moving backwards-in-time, for each population *P_Angola_* and *P_Mozambique_*, population *P_n_* is initialized with a diploid effective population size of *N*_n_ *at Generation Present* and an exponential growth/decay rate of *α*_n=_log(*N*_n_*/N*_n_^’^)*/Generation Split*, where *Generation Split* is defined as the generation at which populations *P_Angola_* and P*_Mozambique_* merge (i.e., lineages can coalesce freely) to form *P_Ancestral_* and *N’_n_* is the diploid effective population size of *P*_n_ at *Generation Split*.

2 *Split with gene flow.* In addition to the parameters defined in (1), moving backwards-in-time, some proportion *Adm_Ang->Moz_* of the lineages in population *P_Mozambique_* migrate to population *P_Angola_* (and vice versa, with proportion *Adm_Moz->Ang_*) at *Generation Gene Flow (GF)*.

To reduce computation time, we additionally include two fixed parameters across both models wherein the merged population *P_Ancestral_* instantaneously changes to a diploid population size of 12,000 (*N_Fixed_*) at generation 7,586. Values were based on the model of human population history presented in Tennessen et al.^[81]^. Using the coalescent simulator *msprime*^[51]^, we generated 10^5^ simulations of chromosome 1 for each scenario defined above with values for each parameter randomly drawn from their corresponding prior distributions (Supplementary Table 9). A mutation rate of 1.25 x 10^-8^ per base-pair per generation was selected and variable recombination rates across the approximately 249 Mb sequence of chromosome 1 were input using inferred genetic distances between sites. Simulated tree sequences were subsequently converted into phased VCF files. To additionally simulate genotyping error-rates associated with low or intermediate coverage sequencing data, we also applied a genotype error rate of 0.001^[52]^ independently for each simulated VCF. Specifically, error was introduced by converting homozygote genotypes to heterozygote and by converting heterozygote genotypes to a randomly chosen homozygote using a custom R script (*github link placeholder*). As our observed data, we used 40 randomly sub-sampled, unrelated Angolan genomes and 40 randomly sub-sampled, unrelated Mozambican genomes. This data was subsequently restricted to approximately 1.5 million biallelic SNPs present on chromosome 1 segregating in these 80 individuals and phased using SHAPEITv2^[78]^. For every simulated and observed VCF we calculate a set of 46 summary statistics as described in Supplementary Table 8 (adapted from REF^[53]^).

We next use the Random Forest ABC model selection procedure from the *abcrf* R-package^[49]^. Applying the *abcrf* function to a combined reference table including all summary statistics with associated simulation parameters and identifying model IDs, we built a model classifier object by growing 1000 decision trees with the same prior probability applied to each model. We also allow *abcrf* to calculate and use the first and second Linear Discriminant functions to build the model classifier. *abcrf* calculates error-rates associated with model classification using an out-of-the bag approach. We ensured that the number of decision trees was sufficient by examining the stability of this error-rate using the *err.abcrf* function. Using this model classifier, we use the *predict* command to estimate the posterior probability of the summary statistics computed using SNP data from Angolans and Mozambicans belonging to either model class.

To estimate posterior distributions and median point estimates of each demographic parameter value (Supplementary Table 9), we use the *neuralnet* method implemented as part of the R-package *abc*^[50]^ with a logit transformation applied to each parameter.

We assess the accuracy of the median points of the posterior distributions by calculating the Mean Absolute Error (MAE), Mean Squared Error (MSE) and Root Mean Squared Error (RMSE) using the *Metrics* R-package (https://github.com/mfrasco/Metrics) by comparing with pseudo-observed parameter values from 1,000 randomly selected simulations (Supplementary Figure 8, Supplementary Table 10). We further ensured posterior distributions captured true uncertainty in parameter estimates by calculating the frequency with which pseudo-observed parameter values associated with 1,000 randomly selected simulations appear within the 2.5 and 97.5 percentile bounds of their corresponding posterior distributions (Supplementary Table 10). We found a priori that 4 neurons in the hidden layer and a 10% tolerance level minimised the average prediction error. Finally, we use our observed summary statistics computed using SNP data from Angolans and Mozambicans alongside the complete set of 100,000 simulations from the *split with gene flow* model to calculate posterior distributions and median point estimates independently for each parameter (Supplementary Table 10).

### 19 Imputation Analyses

As our target datasets for imputing genotypes, we use phased, biallelic, autosomal SNPs from 2,301 self-reported African Americans from the USA collected, sequenced and genotyped as part of the AASP^[55]^ and 1,171 Brazilians collected, sequenced and genotyped as part of SABE^[56]^. To ensure compatibility with reference panels we use UCSC LiftOver^[73]^ alongside the hg38toHg19 chain file (https://hgdownload.cse.ucsc.edu/goldenpath/hg38/liftOver/hg38ToHg19.over.chain.gz) to convert AASP and SABE SNP coordinates from hg38 (https://www.ncbi.nlm.nih.gov/assembly/GCF_000001405.26/) to hg19/GRCh37. We also used 50 Angolans and 10 Mozambicans randomly sub-sampled from our novel collection of whole-genomes as target data. Prior to imputation, SNPs across target datasets were phased using SHAPEITv2^[78]^ and masked at all-but approximately 2.5 million autosomal loci present in the Illumina Omni 2.5 array panel. African ancestry proportions in SABE and the AASP were estimated using ADMIXTURE^[25]^ after merging data from either cohort with genotype data from the 1000G and our newly sequenced samples from Angola and Mozambique and applying the same procedure as outlined in Methods 5. Haplotypes were split into 5Mb chunks and provided in parallel to the IMPUTE2^[82]^ software to impute reference panel genotypes using either the 1000G Panel or the 1000G reference panel merged with unrelated Angolan and Mozambican genomes using the *– merge_reference_panels* command. The 1000G has been shown to perform equal to or better than other, commonly used reference panels (e.g., CAAPA, HRC) when imputing genotypes in African Americans^[55]^. When imputing genotypes in target Angolans and Mozambicans, these 60 target samples were removed from the reference set. Imputed genotypes with an INFO score (r^2^ <0.3) were filtered out. As a metric of imputation accuracy, for each reference panel, we calculate Pearson’s Correlation Coefficient (Dosage R^2^) using imputed genotype dosages across the target dataset and the original, unmasked genotypes as a function of non-reference allele frequency.

## Acknowledgements

This project was partially funded by the Center for Computational, Evolutionary, and Human Genomics, Stanford University, USA (contract no. 60876425). Funding for sequencing of DNA collected from the Cabinda cohort was provided by 23andMe. We thank Dr. Joanna Mountain, Dr. Anjali Shastri, Dr. Steven Micheletti, and Katelyn Kukar for helpful discussions and support of the project. We would like to thank Rosa Kety, Angela Vitor, and Catarina de Jesus from Universidade Onze de Novembro, and Edson Chambala, Stefan Machava, Lourenço Changa and Carlos Dezanove from Universidade Pedagógica for their assistance in sample collection. We are grateful for the administration support of Universidade Onze de Novembro and of Universidade Pedagógica. We would like to acknowledge Professor João Fernando Manuel, Dean of Universidade Onze de Novembro, for discussions about the ethnography of Angola and for his support on this project. The African Genome Variation Project (AGVP) genomic data was obtained from the European Genome-Phenome Archive (ENA) (dataset no. EGAD00001001663). The African American Sequencing Project (AASP) genomic data was obtained from dbGAP (dataset no. phs001102.v1.p1). We would like to thank Professor Michel S. Naslavsky and Dr Marília O. Scliar from the Human Genome and Stem Cell Research Center at the Biosciences Institute of the University of São Paulo, Brazil, for access to the *Saúde Bem Estar e Envelhecimento* project (SABE) genomic data. Finally, we would like to thank the Angolan and Mozambican participants for their invaluable contributions.

