## Supplementary Note for "Whole-genome sequencing of Bantu-speakers from Angola and Mozambique reveals complex dispersal patterns and interactions throughout sub-Saharan Africa"

### DNA Extraction & Sequencing

After obtaining full participant consent, 305 saliva samples were collected from Cabinda, Angola and 196 from Maputo, Mozambique using the Oragene (URLs) collection kit; a strategy previously shown to isolate high levels of endogenous genomic DNA whilst minimizing bacterial DNA content (James et al. 2011). DNA was isolated from each of these 501 saliva samples at the University of Leicester using the following extraction protocol:

DNA was extracted with prepIT•L2P | PT-L2P according to manufacturer's manual protocol from 0.5ml of sample (DNA Genotek) (URLs), with minor deviations. RNA was degraded via introduction of a 30-minute, 37°C incubation with 10mg/ml RNAseA (Thermo Fisher Scientific) (URLs) directly after initial denaturation and DNA release incubation, centrifugation times increased to 5-minutes, ethanol volume increased to 1.5X supernatant volume for DNA precipitation, ethanol wash strength increased to 85% and final resuspension in 10mM Tris HCl pH8.5. Aliquots holding 2-5ug of isolated DNA from 300 Angolan participants were shipped to NGX Bio (URLs) for intermediate-coverage whole genome sequencing after PCR-free library preparation. Of the 196 Mozambican samples, 50 were selected and shipped to Novogene (URLs) for intermediate-coverage, whole-genome sequencing. Reads of length 150 bp were generated by Illumina HiSeq X in both cases, totaling 15,761 gigabases (Gb) of raw sequence data. Paired end reads in FASTQ format were received for all 350 samples (Supplementary Table 1).

### Sample Processing and Quality Control

#### Read Depth

minimap2 v2.11-r797 (sx mode; Li et al. 2018) was used to map FASTQ format-ted paired-end reads generated from each sample against the human reference genome builg hg19/GRCh37 (URLs). Resultant BAM files were sorted according to linear reference coordinates, and duplicate reads were marked with samtools v1.9 mark-dup (Li et al. 2009). Base quality-scores were re-calibrated with GATK v4.0.2.1 BaseRecalibrator and ApplyBQSR (McKenna et al. 2010); using the most recent (as of 9th May 2018) build of the dbSNP150 (URLs) as the set of known variants upon which to build the recalibration/covariation model applied to our sorted BAM files. BAM files were subsequently merged at the individual level, converted to CRAM format and then further compressed with crumble v0.8.1 (Bonfield et al. 2019) using default (compression level 9) parameters.

Read depth statistics were generated from CRAM files using mosdepth v0.2.3 (Pederson and Quinlan 2018) and the sex of each sample was confirmed by comparing the ratio of coverage between the X and Y chromosomes (X/Y ratio). We find two individuals: cab262 and cab188 whose estimated sex was ambiguous or significant outlying compared to the average female or male X/Y ratio (Supplementary Table 1). We take this as evidence of cross-contamination between samples. A further three individuals: cab41, cab191 and cab255 had an estimated sex which disagreed with their given sex. We find no evidence for sample-duplication or contamination in either of these three samples and as such, they were included in all downstream analyses.

After read alignment and duplicate removal, the average autosomal read depth for intermediate-coverage samples was calculated to be 11.56X for Angolans and 12.07X for Mozambicans respectively. Across both sets of individuals, an average of 93.94% +/-2.86 of total reads mapped to the reference genome, validating DNA collection from saliva as a means of collecting high-quality WGS data from human samples. As notable exceptions, two samples: cab91 and cab216 had significantly lower read depth and a higher percentage of unmapped reads than expected, likely due to contamination with pathogenic bacterial DNA during sample collection (see below). The remaining samples had a genome-wide per-site read depth which followed an expected negative-binomial distribution.

### Contamination

To further investigate potential cross-sample/human DNA contamination, we used VerifyBamID2 (Zhang et al. 2018) to calculate ancestry agnostic estimates of contamination fractions per sample (α) using a training set of 100,000 sites from the 1000 Genomes Project Phase 3 (1000 Genomes Project Consortium, 2015). After applying the algorithm to each individual BAM file, we find two samples cab262 (α: 0.32) and cab216 (α: 0.12) which show elevated levels of contamination (previously flagged above). Both were removed from further downstream analyses along with one individual exhibiting extremely low coverage (cab91; Supplementary Table 1). The remaining 347 samples had an estimated contamination level of <1%.

### Unmapped Reads

Recent studies have shown that whole-genome sequencing libraries extracted from saliva samples provide a potentially rich source of information regarding the microbial composition of the oral cavity. In order to assess the origin of potentially contaminating bacterial sequences present in our dataset, we used KrakenUniq (Bretweiser et al. 2018) to perform k-mer based metagenomic classification of reads for which failed to map to the human reference genome. Prior to running KrakenUniq, samtools v1.9 was applied to sample-specific CRAM files to extract high-quality (Q>30) reads for which both mate-pairs were not aligned to the reference build hg19/GRCh37. Reads were then remapped to hg38/GRCh38 (URLs) and an additional 125,715 contigs (296Mb total) recovered as part of an extended African human pan-genome (Sherman et al. 2019). FASTQ files containing all remaining unmapped reads (UMRs) were used as input for KrakenUniq, and taxa-abundance estimates for each sample were generated by comparing sequences with a 31-mer indexed database comprised of all complete microbial genomes deposited in RefSeq (URLs).

As in earlier studies (Samson et al. 2020), we find unmapped reads generated as part of our sequencing efforts represent an abundance of known colonizers of the oral cavity. The most represented genera estimated across each individual sequencing library are Prevotella (mean 5:64 3:5% UMRs) and Haemophilus (mean 5:4 3:2% UMRs) followed by Streptococci (mean 4:5 2:6% UMRs) and Neisseria (mean 4:1 2:2% UMRs), with the remaining UMRs mapping to a further 420 unique microbial genera. Interestingly, we find that the two samples cab91 and cab216 which appeared as significant outliers with regards to the proportion of reads mapped to the human reference sequence (Supplementary Table 1) were also inferred by KrakenUniq to have by far the highest number of non-human reads derived from a single bacterial species. In individual cab91, 305,905,813 (77.23% of total UMRs) reads mapped to *Stenotrophomonus maltophilia* and in cab216 226,791,973 (73.51% of total UMRs) mapped to *Klebsiella varicola* both of which are pathogens known to cause upper-respiratory tract infections in humans (Kanderi et al. 2020, Rodriguez-Medina et al. 2019). Although all precautions were taken prior to this study to avoid sampling from such individuals, these results clearly highlight the fact that sampling saliva in patients who are, for example, suffering from upper-respiratory tract infections is highly unreliable for use in genomic sequencing projects.

### Variant Calling & Filtering

#### Autosomes & X-chromosome

CRAM files were used as input for GATK v4.0.2.1 (McKenna et al. 2010) to jointly call variants across all 347 remaining samples (Supplementary Table 1) following the Best Practices Pipeline (URLs) modified where necessary to fit our intermediate-coverage sample set. Specifically, GVCF files for each sample were generated using the

HaplotypeCaller command. The human reference genome was then split into 348 8Mb contigs (shorter at telomeres) covering autosomes and the X-chromosome and used to parallelize the collection and genotyping of all samples using GenomicsDBImport and GenotypeGVCFs respectively. The resulting set of raw, unfiltered variant calls included 37,048,597 Single-Nucleotide-Polymorphisms (SNPs) (Ts/Tv= 2.00), 9,681,152 short (<150bp) insertions or deletions (INDELs) with 3,903,720 observed multi-allelic sites.

To filter out spurious variant calls e.g., potential sequencing errors or mapping artifacts, we applied the following quality-control procedure (in order):

1. Variants with significant departure from Hardy-Weinberg equilibrium in the direction of excessive heterozygosity (p <3.4e-06, phred-scaled >54.69) using Fisher's exact test (Wigginton et al. 2005) were filtered out. Such variants have been shown to be associated with genotyping error.

2. For SNPs, the Variant Quality-Score Recalibration (VQSR) functionality of GATK v4.0.2.1 was applied with recommended features using the training sets: (1) HapMap 3.3 (truth = true, training = true, known = false, prior 15.0) (2) Omni 2.5 overlapping with 1000G Phase 1 (truth = true, training = true, known = false, prior = 14) (3) 1000G Phase 1 call-set (truth = false, training = true, known = false, prior = 12.0) (4) dbSNP150 (truth = false, training = false, known = true, prior = 2.0) in -mode SNP. A tranche-sensitivity value of 99.5 was selected and variants outside of this cut-off were removed. A tranche sensitivity cut-off of 99.5 was decided after calculating False-Discovery-Rates (FDRs) (estimate of 4% for

chosen cut-off) and True-Positive-Rates (TPRs) (estimate of 94.6% for chosen cut-off) comparing SNP calls in samples also sequenced to 50X coverage.

3. For short INDELs, the VQSR functionality of GATK v4.0.2.1 was applied with recommended features using the training sets: (1) Mills and 1000G gold-standard Indel set (truth = true, training = true, known = false, prior 12.0) (2) Axiom Poly Exome Plus Array set (truth = false, training = true, known = false, prior = 10.0) (3) dbSNP150 (truth = false, training = false, known = true, prior = 2.0) in -mode INDEL. A tranche-sensitivity value of 99 was selected and variants outside of this cut-off were removed. A tranche sensitivity cut-off of 99 was decided after calculating FDRs (estimate of 55% for chosen cut-off) and TPRs (estimate of 70% for chosen cut-off) comparing INDEL calls in samples sequenced to 50X coverage.

4. Variants with >2 variants at a single position (multi-allelic) were removed i.e., only biallelic REF/ALT variants were processed. This includes VCF 4.3 (https://samtools.github.io/hts-specs/VCFv4.3.pdf) formatted spanning deletions (as represented by * in VCF files) as well as overlapping INDEL calls.

5. Variants with >10% missingness across our sample-set were removed.

After filtration, a total of 29,076,696 SNPs (Ts/Tv=2.05) and 3,767,719 INDELs (67% of which were single point changes) present along the autosomes remained along with 885,705 SNPs and 156,559 INDELs on the X-chromosome. Note that uniparental markers (Y-chromosome and mtDNA) were processed separately as outlined below.

It has been previously shown that exploiting correlation in linkage-disequilibrium (LD) between large numbers of individuals alongside genotype uncertainty can improve the quality of genotypes by correcting calls in disagreement with the underlying haplotype (Tom et al. 2017).We calculated posterior genotype probabilities (GP) using Beagle v4.1 (Browning and Browning 2007) with default parameters, using phred-scaled genotype likelihoods (PLs) generated by GATK v4.0.2.1 HaplotypeCaller (McKenna et al. 2010) at 29,962,401 biallelic, genome-wide SNPs and 3,861,191 INDELs which passed our filtration criteria. Male heterozygous (REF/ALT) PLs on the non-pseudo autosomal regions (PAR) of the X-chromosome were truncated to avoid spurious diploid calls. To speed up computation, we parallelized Beagle v4.1 across windows of 25,000 sites with a 1,500-site overlap and subsequently merged generated VCF files. PLINK formatted genetic maps for reference build hg19/GRCh37 used in this processing step were downloaded from the Beagle webpage (URLs). As we find that INDELs across our dataset likely contain a high proportion of false-positives due to the difficulty of calling these variants in intermediate-coverage sequencing data (Fang et al. 2014) we only retain INDELs with a posterior genotype probability >0.9. However, as we find that the number of these short INDELs per sample remains moderately correlated with sequencing coverage, we avoid further interpretation or analyses involving these variants.

Not all positions across the human genome are equal in terms of relative sequence content or uniqueness. Thus, certain regions of the reference genome may be inaccessible to short-read sequencing technologies and therefore exhibit significantly variable genotyping properties. To evaluate study-specific genomic accessibility, we generated a set of “not-passing” bases using a procedure adapted from the 1000G (The 1000 Genomes Project Consortium et al. 2015). Specifically, the base is inferred to be “not-passing” if (1) The reference base is an N. (2) The mean study-wide coverage at that base is less than half the average. (3) The mean study-wide coverage at that base is more than double the average. (4) More than 20% of the reads at that base have a mapping quality (MQ) of 0. After applying the above criteria, we infer a total of 260,731,626 autosomal bases (9% of total) across the autosomes as being “not-passing” or inaccessible based on a study-wide autosomal average read depth of 4461X calculated using mosdepth v0.2.3 (Pederson and Quinlan 2018) and a further 9,131,659 bases (6% of total) across the X-chromosome (evaluated separately due to haploid males) based on an average read depth of 3400X. Across the entire call set, we find only 50,607 (0.2%) called SNPs fall outside of our accessible genome mask, suggesting our variant calling and filtration procedure was reasonably effective in removing spurious variant calls within these regions. Although we do not explicitly filter out these loci from our release, we do exclude these variants from all downstream population genetic analyses, as such regions are likely to be enriched for false positives.

To further validate our variant-calling procedure, SNP density across the autosomes was computed using VCFTOOLS v1.014 (Danacek et al. 2011) using non-overlapping bins of 100kb. As expected, we find low SNP density in centromeric and telomeric regions and high density in known mutational hotspots such as the Major Histocompatibility Complex (MHC) on chromosome 6 (data not shown). When comparing the normalized SNP density observed in our dataset to that of the 1000G (Pearson’s $r^2$ = 0.92) or the

AGVP (Pearson's $r^2 = 0.97$) we find a high correspondence. The lower correlation observed with the 1000G is likely due to variant calling being performed across a larger, more cosmopolitan set of samples relative to this study and the AGVP (Supplementary Table 2).

Y-Chromosome

As in Poznik et al. 2013, we first define accessible regions of the Y-chromosome using 184 male-specific CRAM files present in our study by emitting sites whereby (1) Read-depth deviated from a study-wide exponentially weighted moving average (EWMA) estimated across contiguous 1kb intervals (2) More than 10% of the reads at that site had a MQ of 0.

After applying the above procedure, we called SNPs across 11.6 MB of accessible Y-sequence using GATK v3.8 Unified Genotyper (McKenna et al. 2010) with additional flags –stand_call_conf 30 -mbq 30 -gt mode DISCOVERY. Heterozygous genotype calls were set to missing and we subsequently filtered out sites with greater than 10% missingness across the sample set. The median accessible study-wise read depth was 1259X (as calculated by mosdepth v0.2.3) and sites with a depth of more than three median absolute deviations above or below this value were excluded along with multi-allelic sites. A total of biallelic 13,091 SNPs remained, with 3,352 included in the most recent (as of 5th July 2020) International Society of Genetic Genealogy (ISOGG) database (URLs). We used Yhaplo (Poznik et al. 2016) to assign haplogroups to each sample and compared this with haplogroups called in 1,233 males from the 1000G, estimated using the same procedure.

Mitochondrial DNA

mtDNA sequences from 338 high-quality, unrelated samples were processed using a consensus calling strategy like the one employed in Li et al. 2015. Specifically, reads not mapped to the nuclear genome (hg19/GRCh37) were remapped to the Revised Cambridge Reference Sequence (URLs) to a mean depth of 1136X. Consensus sequences were called using the majority rule, requiring minimum base quality of 30 with N (no call) generated if the alternative allele frequency was greater than 0.3. Resultant whole mtDNA FASTA files were subsequently aligned along with 2,534 sequences from the 1000G using MAFFT v7.222 (Katoh et al. 2002). Haplogrep2 v2.1.1 (Weissensteiner et al. 2016) was used to predict haplogroups with PhyloTree build 17 (van Oven and Kayser 2009) and maximum-likelihood based phylogenetic trees were constructed using FastTree v2.1 (Price et al. 2010) with flags -nt -gtr -spr 4.

Merging with external data

Defining Sample Origin

Estimates for the number of Bantu languages across sub-Saharan Africa range from between 440 (Guthrie) and 680 often dividing regions both geographically and culturally. Ethnologue (URLs) notes 44 and 42 extant Bantu languages in Angola and Mozambique respectively, with many regional dialects only beginning to be described in detail (Boeston et al. 2012, https://www.yumpu.com/en/document/read/17832108/kikongo-dialect-continuum-internal-and-external-classification-llacan). To assign ancestry labels to each study participant, we follow a stepwise process whereby individuals are described using (1) their country of origin (Angola, Mozambique) (2) broad language/ethnic group (e.g., Kongo, Ovimbundu, Makua, Tswa-Ronga) (3) self-reported primary language/dialect of both maternal and paternal parents and grandparents if available (e.g., Ibinda, Kiyombe, Chuwabu, Tswa). Individuals with mixed broad parental languages (e.g., Kikongo/Kimbundu) were excluded from the HOA dataset merging to aid in interpretability. Each individual's population label in our merged dataset can be found in Supplementary Table 1.

### Merging procedure

Throughout the study, we analyse our novel collection of samples in the context of autosomal genotype data generated by earlier studies (The 1000 Genomes Project Consortium et al. 2015, Gurdasasani et al. 2015, Lazaridis et al. 2014, Lazaridis et al. 2016, Llorente et al. 2015, Skoglund et al. 2017, Prendergast et al. 2019, Schlebusch et al. 2017, Lopez et al. 2020, Lipson et al. 2020, Wang et al. 2020, Fan et al. 2019, Mallick et al. 2016). Each curated dataset is summarised in Supplementary Table 2 (WGS) and Supplementary Table 6 (HOA) including given population labels. Usage of each merged dataset is dependent on the question of interest and specified in the main text, when necessary, alongside any further analysis-specific filtration of sites.

For each dataset, prior to merging individuals across studies each REF/ALT allele was checked against the reference build hg19/GRCh37 using bcftools v1.9 norm –check-ref (Li 2009), flipped when necessary and subsequently annotated using a hexadecimal 64-bit VariantKey (described in Asuni and Wilder 2019) encoding CHR/POS/REF/ALT to ensure maximum compatibility between datasets. A/T and C/G alleles were removed to mitigate strand ambiguity and SNPs at shared coordinates were merged using bcftools v1.9 merge. To reduce the effects of spurious genotype calls on downstream population genetic analyses, SNPs in difficult to sequence regions were filtered using a merged BED file (generated using bedtools v2.28.0 (Quinlan and Hall 2010) multiinter) including: (1) Our accessible genome mask (see above). (2) Heng Li's Low-Complexity Region (LCR) mask (URLs). (3) The hg19 ENCODE blacklist (URLs). (4) Segmental duplications from UCSC (URLs). We also filtered out one of every pair of related samples (< fourth degree) using the same process using KING as described above.

High-coverage ancient genomes

BAM files from three high-coverage ancient samples: a 2,000 year old individual from South Africa (Ballito Bay A; baa001) as described in Schlebusch et al. 2017, a 4,500 year old individual from Ethiopia (Mota; GB20) as described in Llorente et al. 2015 and an 8,000 year old individual from Cameroon (Shum Laka; I10871) as described in Lipson et al. 2020 were downloaded, processed and subject to diploid variant calling using the procedure outlined in Schlebusch et al. 2017, Supplementary Materials 4.1. Specifically, for each BAM file, we set base quality scores (BQ) of Ts in the first 5bp of each read and As in the last 5bp of each read to 2. GATK v3.8 (McKenna et al. 2010) was used to realign INDELs with the 1000G callset used as a reference (1000 Genomes Project Consortium, 2015). GATK Unified Genotyper (McKenna et al. 2010) was subsequently used to call diploid genotypes with the parameters *-stand_call_conf 50.0 –mbq 40 –contamination 0.02 -out_mode EMIT_ALL_SITES -gt_mode GENOTYPE GIVEN ALLELES* with variants within the dbSNP142 (URLs) given as known sites using the *–alleles* flag. Calls flagged as "Low Quality" were subsequently filtered out.

Chimpanzee reference genome

As a representative outgroup to all human populations, we use haploid genotypes from the Chimpanzee reference genome (URLs) aligned to the human reference hg19/GRCh37 (GRCh37). axtNet alignment files were downloaded from the UCSC (Chimp Human Alignment) and used to generate VCF files using a custom python script (available upon request).

### URLs

Oragene Saliva Kit (DNA Genotek): https://www.dnagenotek.com/ROW/products/collection-human/oragene-dna/500-series/OG-500.html

Oragene prepIT (DNA Genotek): https://www.dnagenotek.com/US/products/reagents-preparation/prepIT/PT-L2P.html

RNAseA (Thermo Fisher): https://www.thermofisher.com/order/catalog/product/EN0531

NGX Bio: https://ngxbio.com/

Novogene: https://en.novogene.com/

hg19/GRCh37: https://www.ncbi.nlm.nih.gov/assembly/GCF_000001405.13/

hg38/GRCh38: https://www.ncbi.nlm.nih.gov/assembly/GCF_000001405.26/

panTro5: https://www-ncbi-nlm-nih-gov.ezproxy3.lib.le.ac.uk/assembly/GCF_000001515.7/

rCRS: https://www.mitomap.org/MITOMAP/HumanMitoSeq

dbSNP150: ftp.ncbi.nih.gov/snp/organisms/human_9606/VCF/

RefSeq: https://www-ncbi-nlm-nih-gov.ezproxy3.lib.le.ac.uk/refseq/

ISOGG: https://isogg.org/tree/ISOGG_YDNA_SNP_Index.html

GATK Best Practices Pipeline:

https://www-ncbi-nlm-nih-gov.ezproxy3.lib.le.ac.uk/refseq/

Beagle Genetic Map: http://bochet.gcc.biostat.washington.edu/beagle/genetic_maps/

1000 Genomes Genetic Map: https://github.com/joepickrell/1000-genomes-genetic-maps

Ethnologue : https://www.ethnologue.com/

Low Complexity Regions Mask: https://github.com/lh3/varcmp/raw/master/scripts/LCR-hs37d5.bed.gz

ENCODE Blacklist: https://www.encodeproject.org/annotations/ENCSR636HFF/

UCSC Segmental Duplications:

https://humanparalogy.gs.washington.edu/build37/build37.htm

Chromosome Painting: http://paintmychromosomes.com/
