## Supplementary Figure for "Whole-genome sequencing of Bantu-speakers from Angola and Mozambique reveals complex dispersal patterns and interactions throughout sub-Saharan Africa"

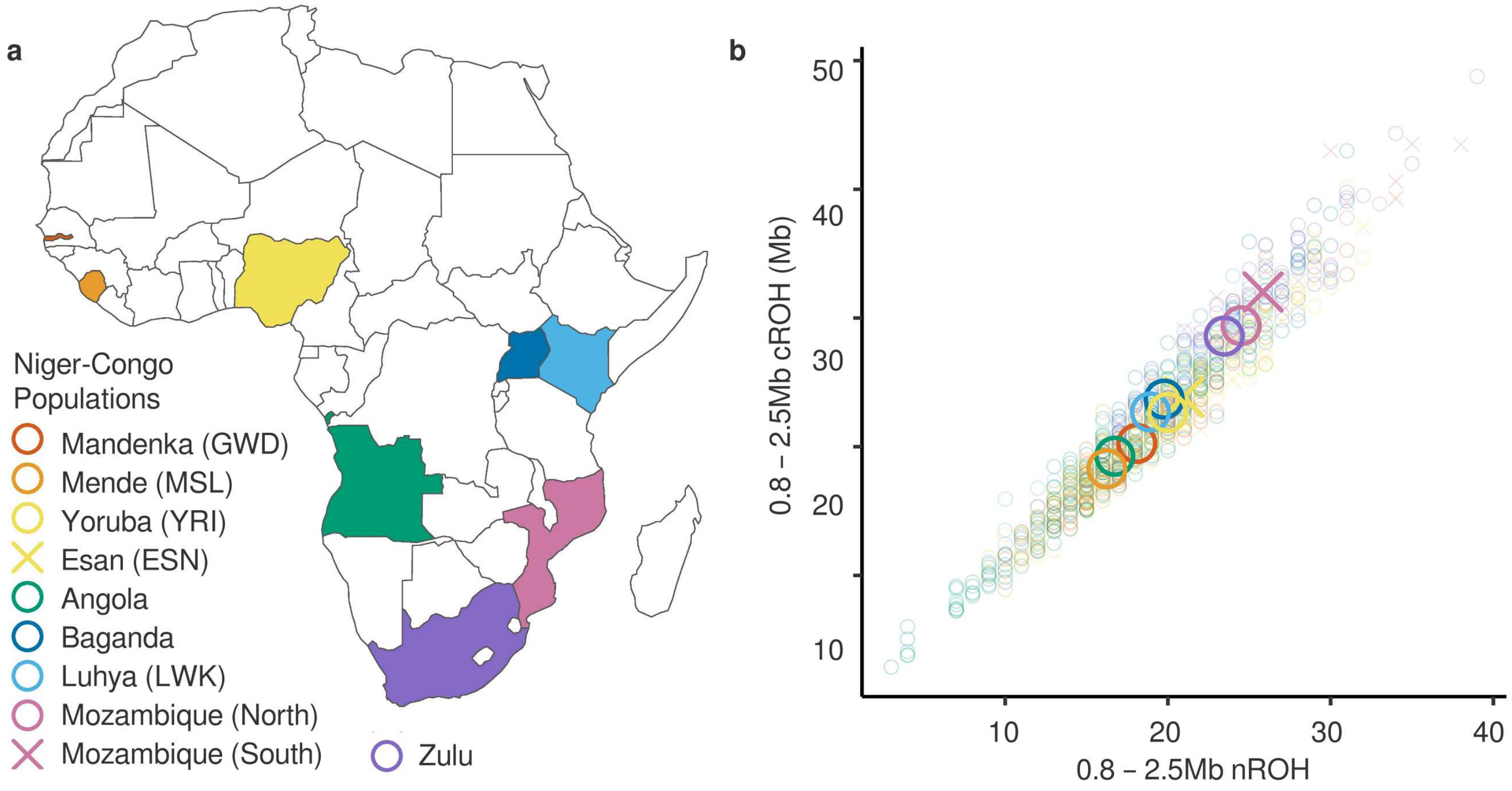


**Supplementary Figure 1.** (**a**) Legend denoting colour and shape of points corresponding to an individual from Niger-Congo populations present in a merged dataset consisting of newly sequenced Angolans and Mozambicans, the 1000G and the AGVP (**b**) Average number of short Runs-Of-Homozygosity (nROH) and cumulative short ROH (cROH) per population in the merged dataset. It should be noted that recent admixture with autochthonous populations in the Luhya (LWK), Baganda, and Zulu is likely to reduce ROH.

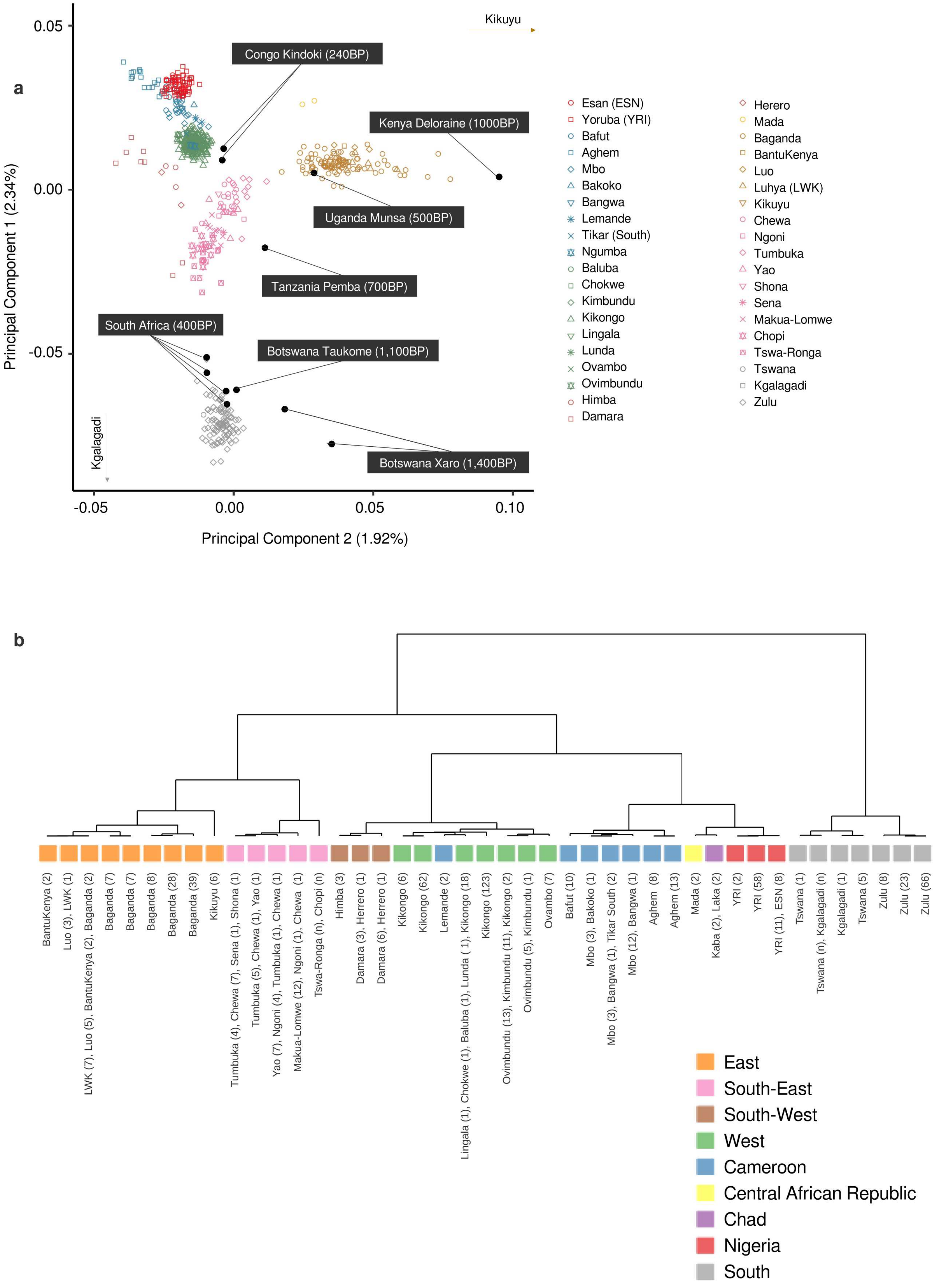

a
Kikuyu
Congo Kindoki (240BP)
Kenya Deloraine (1000BP)
Uganda Munsa (500BP)
Tanzania Pemba (700BP)
South Africa (400BP)
Botswana Taukome (1,100BP)
Botswana Xaro (1,400BP)
Kgalagadi
0.05
0.00
-0.05
0.10
Principal Component 1 (2.34%)
Principal Component 2 (1.92%)
Esan (ESN)
Yoruba (YRI)
Bafut
Aghem
Mbo
Bakoko
Bangwa
Lemande
Tikar (South)
Ngumba
Baluba
Chokwe
Kimbundu
Kikongo
Lingala
Lunda
Ovambo
Ovimbundu
Himba
Damara
Herero
Mada
Baganda
BantuKenya
Luo
Luhya (LWK)
Kikuyu
Chewa
Ngoni
Tumbuka
Yao
Shona
Sena
Makua-Lomwe
Chopi
Tswa-Ronga
Tswana
Kgalagadi
Zulu
b
BantuKenya (2)
Luo (3), LWK (1)
LWK (7), Luo (5), BantuKenya (2), Baganda (2)
Baganda (7)
Baganda (7)
Baganda (8)
Baganda (28)
Baganda (39)
Kikuyu (6)
Tumbuka (4), Chewa (7), Sena (1), Shona (1)
Tumbuka (5), Chewa (1), Yao (1)
Yao (7), Ngoni (4), Tumbuka (1), Chewa (1)
Makua-Lomwe (12), Ngoni (1), Chewa (1)
Tswa-Ronga (n), Chopi (n)
Himba (3)
Damara (3), Herrero (1)
Damara (6), Herrero (1)
Kikongo (6)
Kikongo (62)
Lemande (2)
Lingala (1), Chokwe (1), Baluba (1), Lunda (1), Kikongo (18)
Kikongo (123)
Ovimbundu (13), Kimbundu (11), Kikongo (2)
Ovimbundu (5), Kimbundu (1)
Ovambo (7)
Bafut (10)
Mbo (3), Bakoko (1)
Mbo (3), Bangwa (1), Tikar South (2)
Mbo (12), Bangwa (1)
Aghem (8)
Aghem (13)
Mada (2)
Kaba (2), Laka (2)
YRI (2)
YRI (58)
YRI (11), ESN (8)
Tswana (1)
Tswana (n), Kgalagadi (n)
Kgalagadi (1)
Tswana (5)
Zulu (8)
Zulu (23)
Zulu (66)
East
South-East
South-West
West
Cameroon
Central African Republic
Chad
Nigeria
South

**Supplementary Figure 2.** Population structure of Angolans and Mozambicans in the context of Bantu-speaking groups (and closely related Nigerians) present in the HOA dataset. (**a**) Principal Components Analysis with eigenvectors (PC1 and PC2) constructed using modern populations (shown in legend) and ancient Bantu-associated genomes projected onto these axes. (**b**) fineSTRUCTURE inferred dendrogram relating 43 clusters calculated using the chunk counts matrix generated from CHROMOPAINTER using the *"all-Bantu-copying"* model population configuration (see Methods 12). Note that this does not include ancient Bantu-related individuals due to the prerequisite of phased, diploid genotypes.

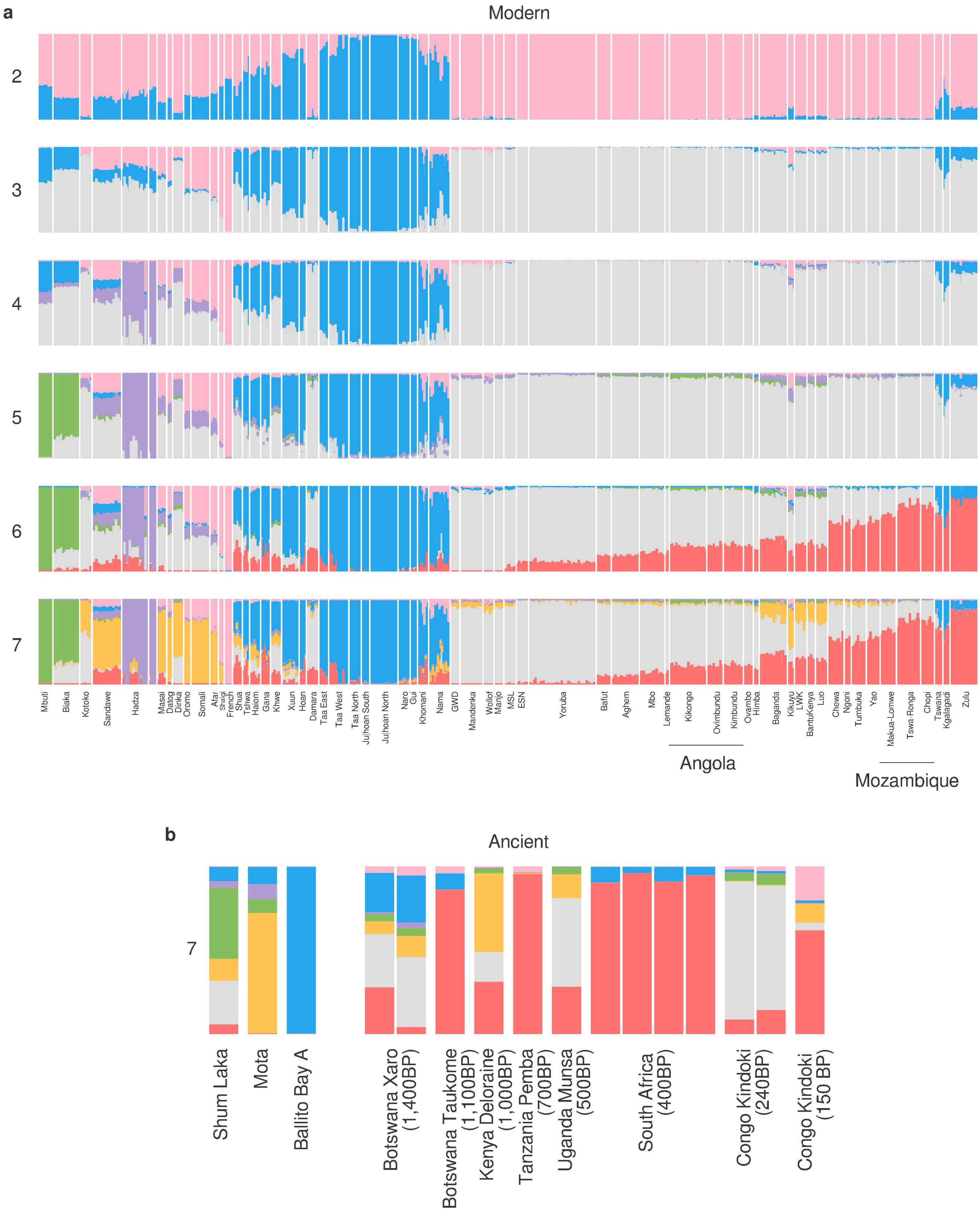
a
Modern
2
3
4
5
6
7
Mbuti
Biaka
Kotoko
Sandawe
Hadza
Masai
Datog
Dinka
Oromo
Somali
Afar
Shaigi
French
Shua
Tshwa
Haiom
Gana
Khwe
Xuun
Hoan
Damara
Taa East
Taa West
Taa North
Ju|hoan South
Ju|hoan North
Naro
Gui
Khomani
Nama
GWD
Mandenka
Wollof
Manjo
MSL
ESN
Yoruba
Bafut
Aghem
Mbo
Lemande
Kikongo
Ovimbundu
Kimbundu
Ovambo
Himba
Baganda
Kikuyu
LWK
BantuKenya
Luo
Chewa
Ngoni
Tumbuka
Yao
Makua-Lomwe
Tswa-Ronga
Chopi
Tswana
Kgalagadi
Zulu
Angola
Mozambique
b
Ancient
7
Shum Laka
Mota
Ballito Bay A
Botswana Xaro (1,400BP)
Botswana Taukome (1,100BP)
Kenya Deloraine (1,000BP)
Tanzania Pemba (700BP)
Uganda Munsa (500BP)
South Africa (400BP)
Congo Kindoki (240BP)
Congo Kindoki (150 BP)

**Supplementary Figure 3.** Unsupervised ADMIXTURE clustering of Angolans and Mozambicans in a pan-African context. (**a**) Modern populations at $K$=2 to $K$=6 (**b**) Ancient populations, including early hunter-gather populations from Cameroon (Shum Laka, 8,000 years old), Ethiopia (Mota, 4,000 years old) and South Africa (Ballito Bay, 2,000 years old) and Bantu-associated individuals whose DNA was extracted from remains present at sites from 1,400 to 120 years ago at $K$=6.

6 represents the best-guess at matching $K$ to the number of true ancestral populations (lowest cross-validation error). However, such interpretations are likely to be affected by population-specific drift and unreasonable assumptions regarding the divergence history of populations across Africa. For example, the Mbuti here appear entirely unadmixed, but fastGLOBETROTTER results clearly shows recent admixture has occurred in the history of this group (Figure 2, Supplementary Table 7).

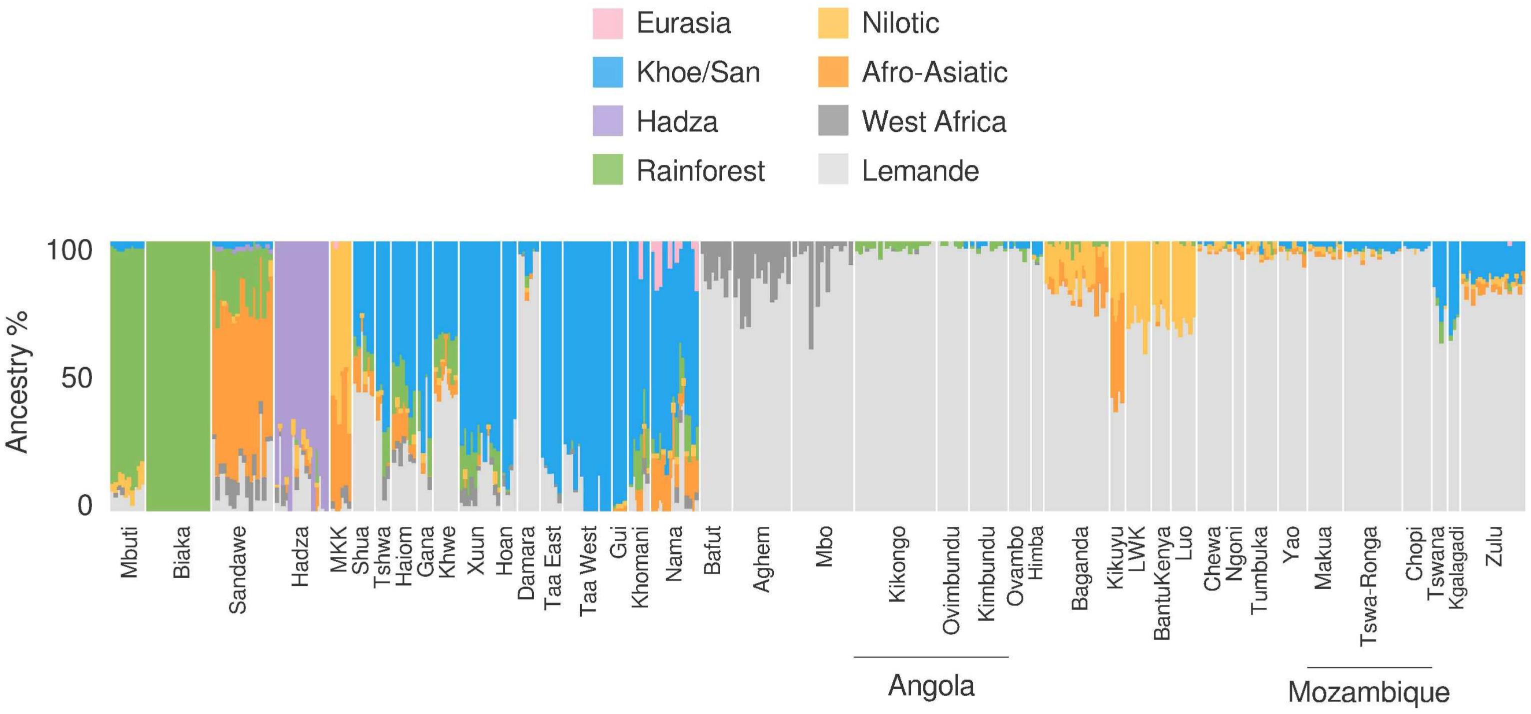


**Supplementary Figure 4.** SOURCEFIND inferred ancestry compositions of Angolans and Mozambicans in a pan-African context performed using a matrix describing the total length of pairwise shared haplotype chunks lgenerated using the *"no-Bantu-copying model"* (see Methods 12). In this model, Bantu-speakers from across sub-Saharan Africa (see "Bantu" super-population in Supplementary Table 6) were excluded as possible donors to CHROMOPAINTER, other than the Lemande. This is to highlight the fraction of additional, autochthonous non-Bantu-related (Lemande) ancestry present in Angolans and Mozambicans when compared to neighbouring groups that may have occurred e.g. through later interactions and admixture with local groups.

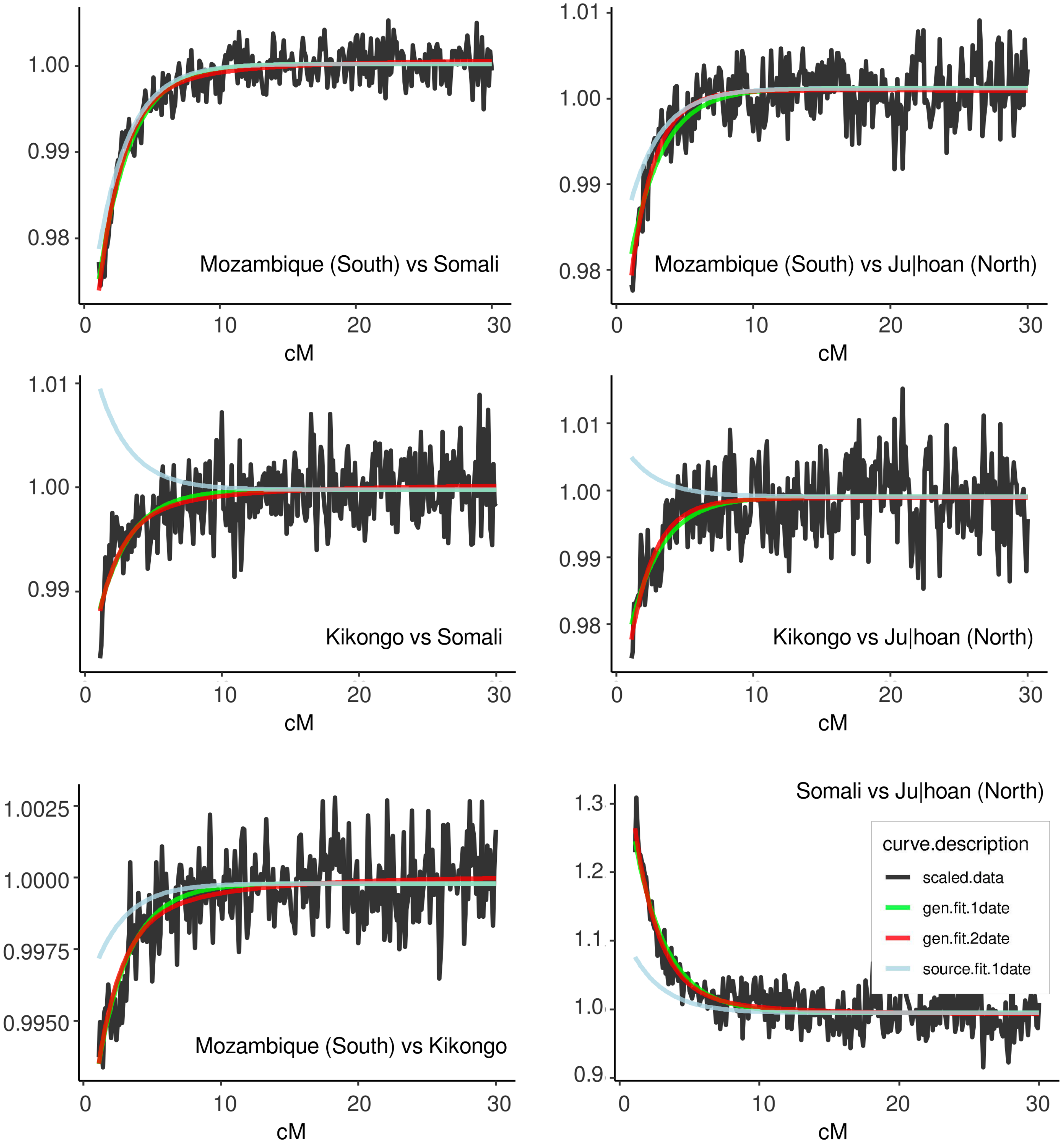


**Supplementary Figure 5.** fastGLOBETROTTER coancestry curves showing evidence of complex multiway admixture in a combined set of Malawians and North Mozambicans (see Supplementary Table 7). Inference of multi-way admixture is also observed when analysing these groups by population label (Yao, Tumbuka, Chewa, Ngoni, Mozambique (North)) as shown in Figure 2 of the main text. Multi-way admixture is inferred when all pairwise combinations of three source groups (here, Mozambique (South), Kikongo, Somali or Julhoan (North)) generate curves showing haplotype length increasing with genetic distance.

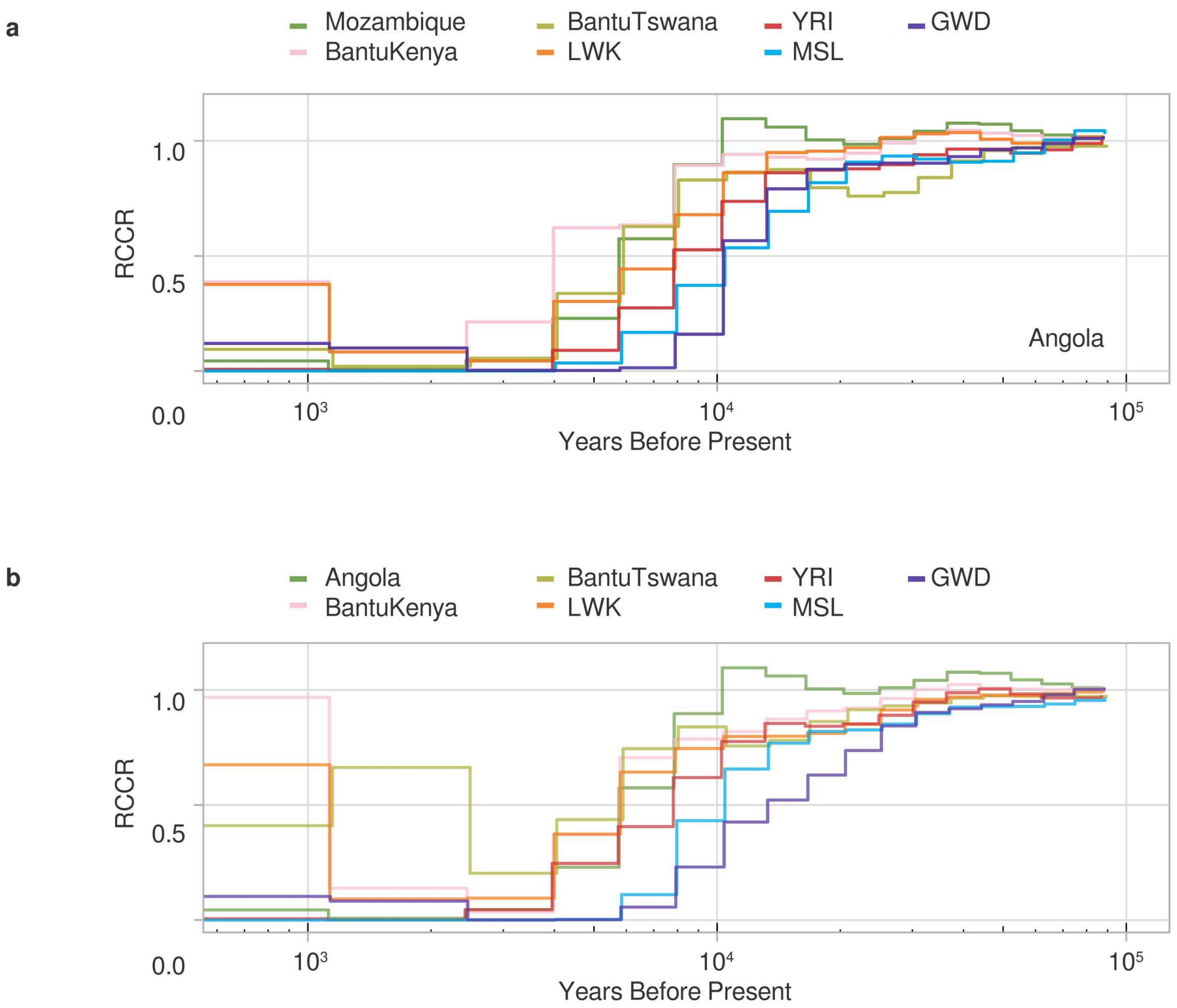


**Supplementary Figure 6.** (**a**) Separation history of Angolan (Kikongo) estimated MSMC2 with two-high coverage (50X) whole-genomes from Angolan and two from either Mozambicans (Tswa-Ronga) or Niger-Congo speaking populations from the Simmons Genome Diversity Project. Separation times were taken as the first generation going backwards-in-time in which RCCR is greater than or equal to 0.5 (**b**) As in a but for Mozambican (Tswa-Ronga).

LWK, Luhya from Kenya; YRI, Yoruba from Nigeria; MSL, Mende from Sierra Leone; GWD, Mandenka from The Gambia; BantuKenya, Bantu-speakers from Kenya (HGDP); BantuTswana, Tswana-speakers from South Africa.

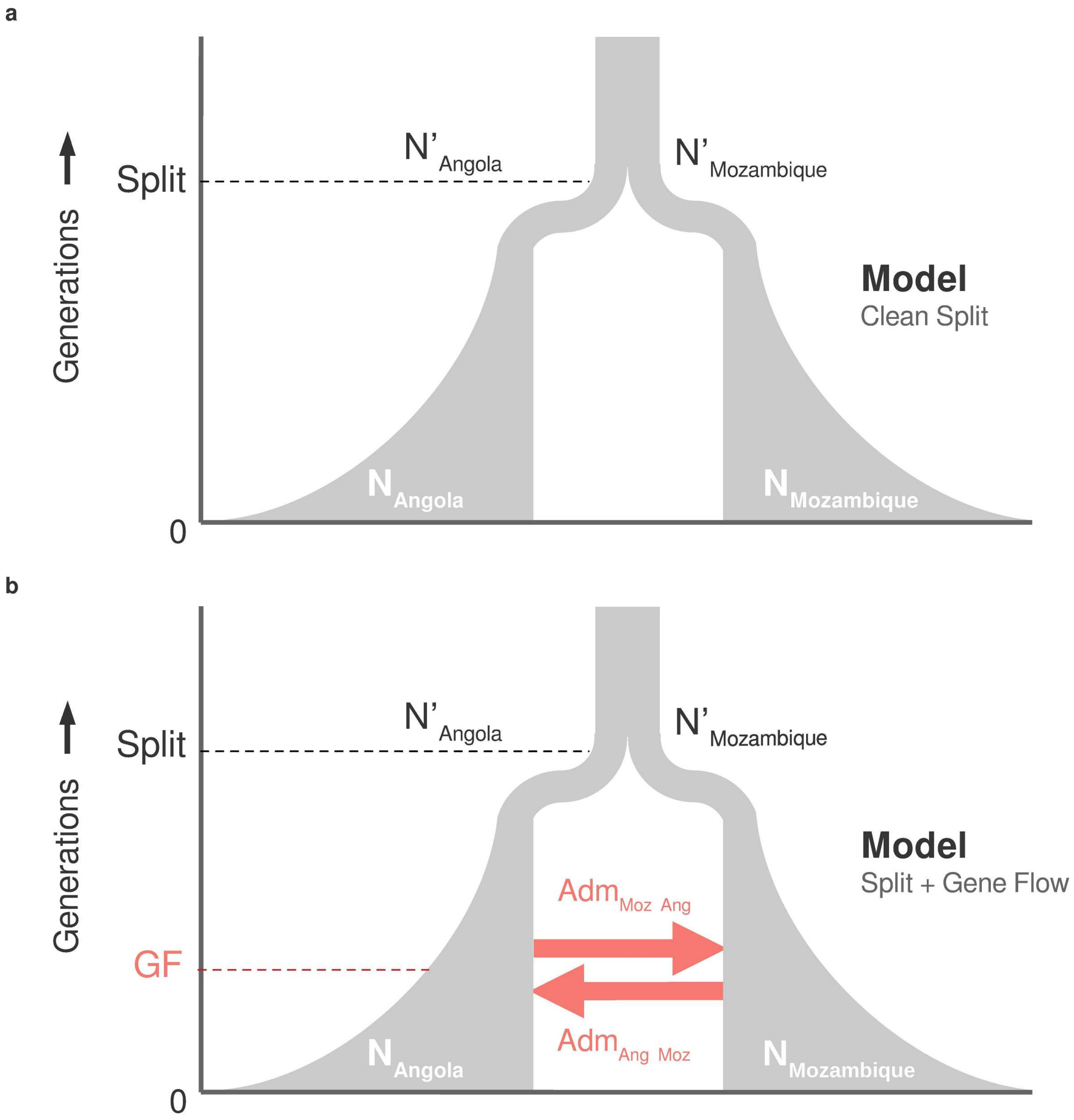


**Supplementary Figure 7.** Models of population separation between the ancestors of Bantu-speakers from Angola and Mozambique. (**a**) *Clean Split* model includes as parameters: the divergence time generation Split and the population sizes $N'_{Mozambique}$ and $N'_{Angola}$ at time generation Split which grow/decay exponentially until generation 0 until size $N_{Mozambique}$ and $N_{Angola}$ respectively as shown in grey. (**b**) *Split with Gene Flow* model additionally includes the parameters $Adm_{Moz\text{->}Ang}$ and $Adm_{Ang\text{->}Moz}$ which describe some proportion of lineages migrating between populations at generation Gene Flow (GF) as shown in red.

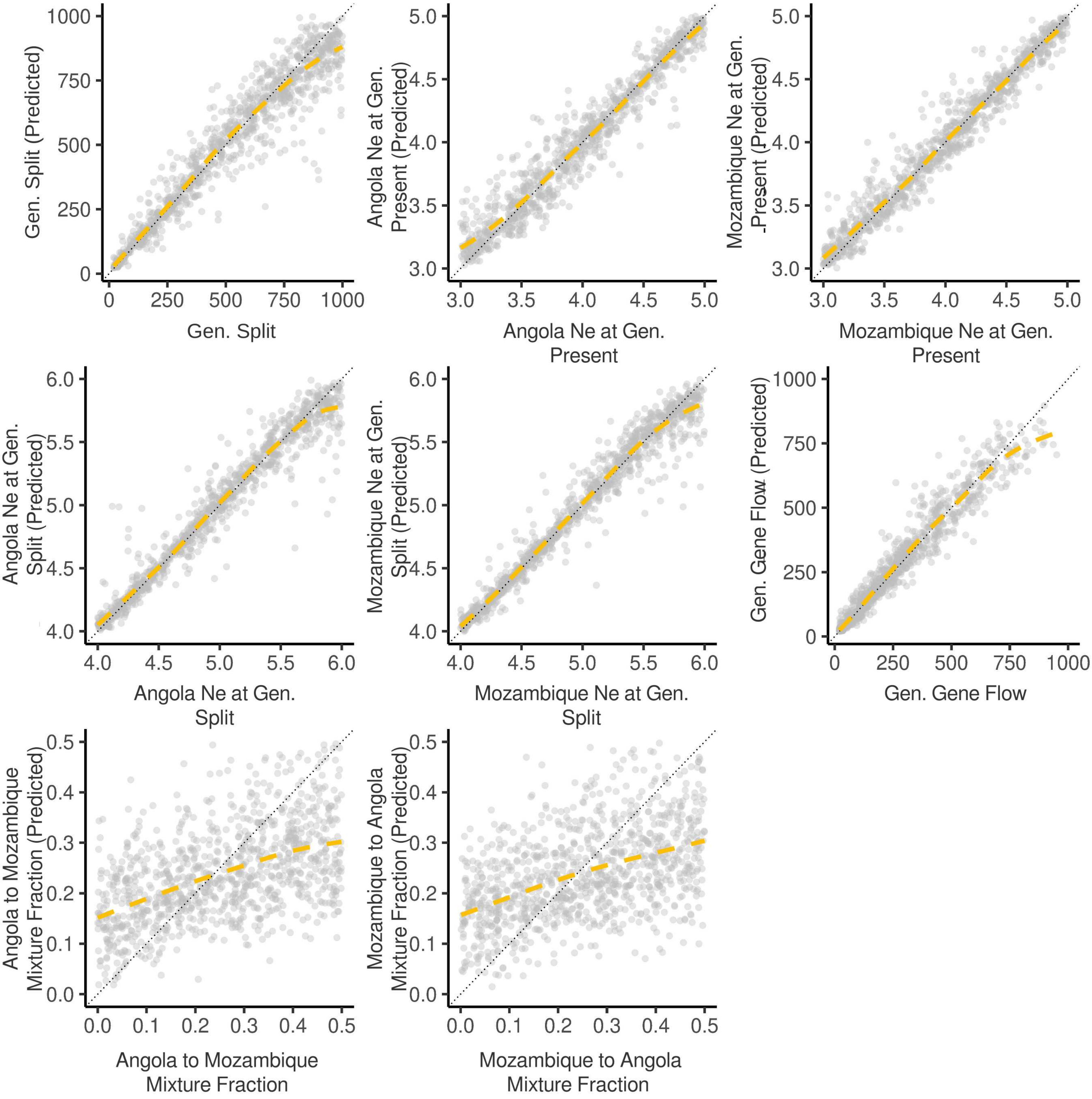


**Supplementary Figure 8.** Accuracy of *abc* ('neuralnet') at estimating the true parameter values using reference tables of summary statistics generated under the *split with gene flow* demographic model (Supplementary Figure 7). For 1,000 pseudo-observed simulations from which the true values of the parameters were known, for each parameter value, *abc* was used to estimate the posterior distribution of the parameter, and the median of the distribution was taken as our estimate of the true parameter value. Yellow line is the average of the median point estimates, dotted line is x=y.

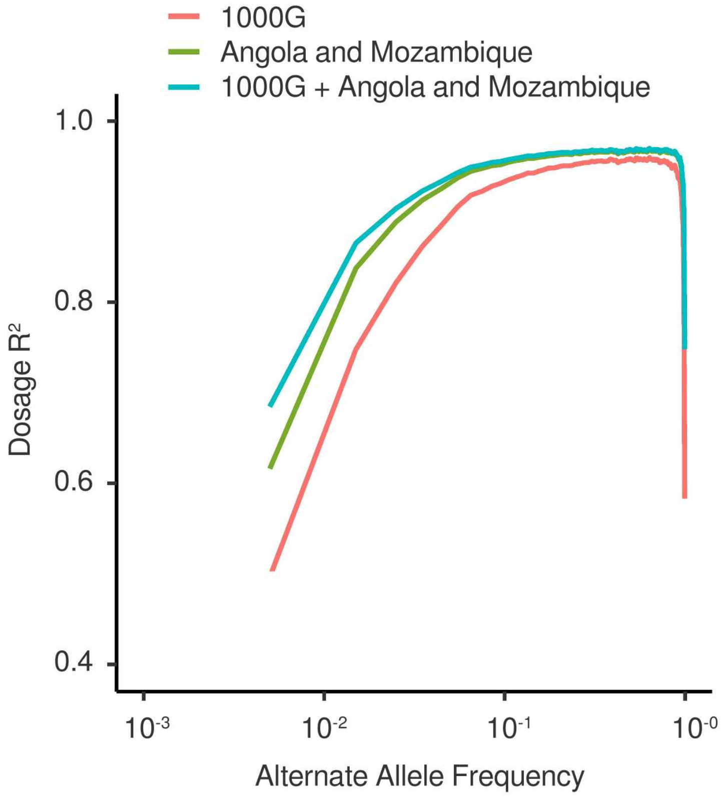


**Supplementary Figure 9.** Dosage R2 (Pearson's squared correlation coefficient) of called genotype vs genotypes imputed into 10 Mozambicans and 50 Angolans using either the 1000G reference panel, the remaining newly sequenced genomes from Angola and Mozambique, or a merged reference panel including the 1000G and newly sequenced genomes from Angola and Mozambique as a function of alternate allele frequency.

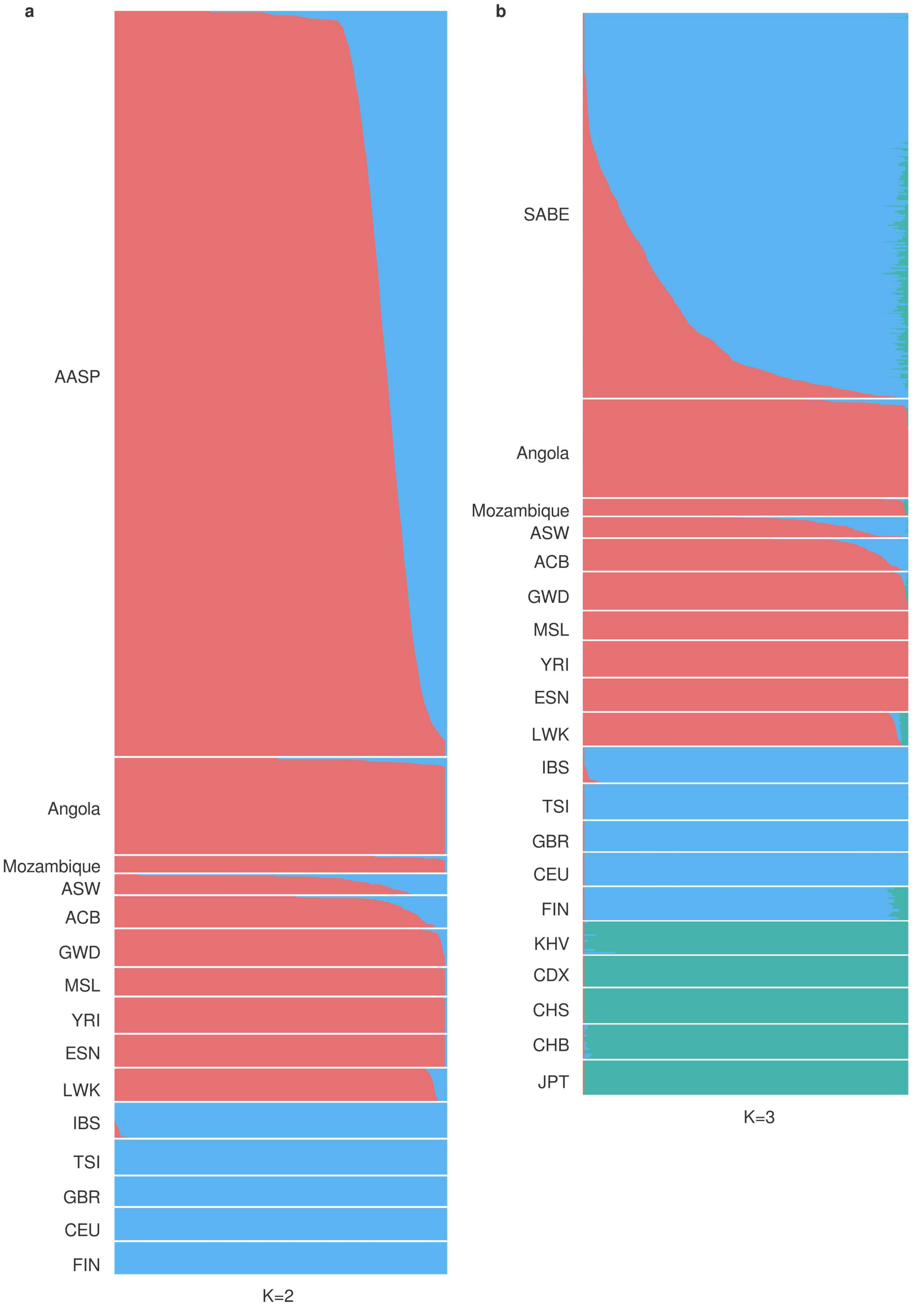
a
b
AASP
Angola
Mozambique
ASW
ACB
GWD
MSL
YRI
ESN
LWK
IBS
TSI
GBR
CEU
FIN
K=2
SABE
Angola
Mozambique
ASW
ACB
GWD
MSL
YRI
ESN
LWK
IBS
TSI
GBR
CEU
FIN
KHV
CDX
CHS
CHB
JPT
K=3

**Supplementary Figure 10.** (**a**) ADMIXTURE clustering at K=2 showing the proportion of West African-related (red) and European-related (blue) ancestry in individuals from the African American Sequencing Project (AASP) alongside samples from Angola, Mozambique, 1000G-AFR, and 1000G-EUR. (**b**) ADMIXTURE clustering at K=3 showing the proportion of West African-related (red), European-related (blue), and East Asian (also a proxy for Native Amerindian) (blue-green) ancestry in individuals from the Saúde Bem Estar e Envelhecimento project (SABE) project alongside samples from Angola, Mozambique, 1000G-AFR, 1000G-EUR, and 1000G-EAS.
